# Trace metals availability controls terminal electron acceptor utilization in *Escherichia coli*

**DOI:** 10.1101/2025.01.08.631794

**Authors:** Annarita Ricciardelli, Benoit de Pins, Jacopo Brusca, Monica Correggia, Luciano Di Iorio, Martina Cascone, Marco Giardina, Stefany Castaldi, Rachele Isticato, Roberta Iacono, Marco Moracci, Nunzia Nappi, Antonino Pollio, Costantino Vetriani, Serena Leone, Angelina Cordone, Donato Giovannelli

**Author notes:** These authors equally contributed to this work (Annarita Ricciardelli, Benoit de Pins).

## Abstract

Trace metals play an essential role in the metabolism of all living organisms and many metal-containing enzymes contribute to key physiological and ecological processes such as aerobic and anaerobic respiration, photosynthesis, carbon and nitrogen fixation. Despite this, trace metals’ potential to control microbial functional diversity and metabolic shifts is unknown. Here we demonstrate that the availability of trace metals controls electron acceptors utilization in *Escherichia coli*. Physiological and proteomic data show that trace metals depleted cultures have significantly reduced growth, start fermentation and increase energy expenditure for metal homeostasis even when more energetically favourable electron acceptors are present. Overall these results suggest how evolutionary and competitive pressures arising from changes in biological trace metals availability in deep time have contributed to shaping evolution and competition.

## Main Text

All life relies on redox chemistry for its existence (*1, 2*). Thermodynamically favourable redox reactions are used by all living organisms to derive energy required for cell maintenance and growth. These reactions are catalysed by specialised enzymes, called oxidoreductases, which control electron flow from electron donors to final electron acceptors, contributing to sustaining the thermodynamic disequilibrium of natural systems (*1*). Several oxidoreductases, particularly those functioning at the cell-environment interface (biogeochemical oxidoreductases, *sensu* (*2*)), rely on metal cofactors that are finely tuned to the redox potential of their substrates (*3*). The diversity of metals used as key catalytic centers by these enzymes remains to be explored (*4*–*6*). Most metal-containing oxidoreductases use transition metals—mainly Fe, Cu, Co, Ni, W, Mo, V, and Mn—in their structure, either alone or as part of complex organometallic cofactors (*7*). Despite their low abundance in the environment, trace metals might play a key role in influencing microbial functional diversity and in shaping microbial community composition (*4*). For example, Fe has been shown to be one of the primary limiting nutrients for primary productivity in the ocean, and competition for iron under iron-limiting conditions may drive community structure and ultimately evolution (*8*–*10*).

Environmental studies suggest that microorganisms in natural populations are capable of adapting and competing in fluctuating environments using alternative electron acceptors (*2, 4, 11*). To verify how widespread the ability to use alternative electron acceptors is in microorganisms, we have annotated ∼52,500 metagenome-assembled genomes (MAGs) from the genomic catalog of Earth’s microbiomes (*12*) that includes MAGs from soils, oceanic water column, marine sediments, subsurface ecosystems, human gut and built environments (Table S1) for genes encoding six main terminal reductases (Table S2). The results reveal that ∼78.1 % of the MAGs encode for more than one terminal oxidoreductase, conferring to the microorganisms who encode them the ability to use alternative electron acceptors (Fig. 1). These include terminal reductases involved in oxygen respiration (both aerobic and microaerophilic), nitrate reduction, sulfate reduction, and thiosulfate reduction. This diversity of oxidoreductases relies on diverse metal cofactors (*2*). For example, oxygen reductases depend on either copper or iron for aerobic and microaerophilic respiration, respectively, while nitrate reductases require molybdenum in their catalytic core. Our analysis reveals that of the 40,283 MAGs for which at least one terminal reductase was identified, ∼37.9 % of the MAGs can use copper, molybdenum and/or iron as cofactors for their terminal reductases, while only ∼29.4 % relies on a single trace metal as cofactor (Fig. 1). Overall, ∼70.6 % of the MAGs annotated relies on diverse metal cofactors for its terminal reductases. This suggests that microorganisms in natural ecosystems need a variety of metals to access diverse terminal electron acceptors, and that limited metal availability may constrain the use of specific metabolic pathways (*4*).

**Figure 1.**
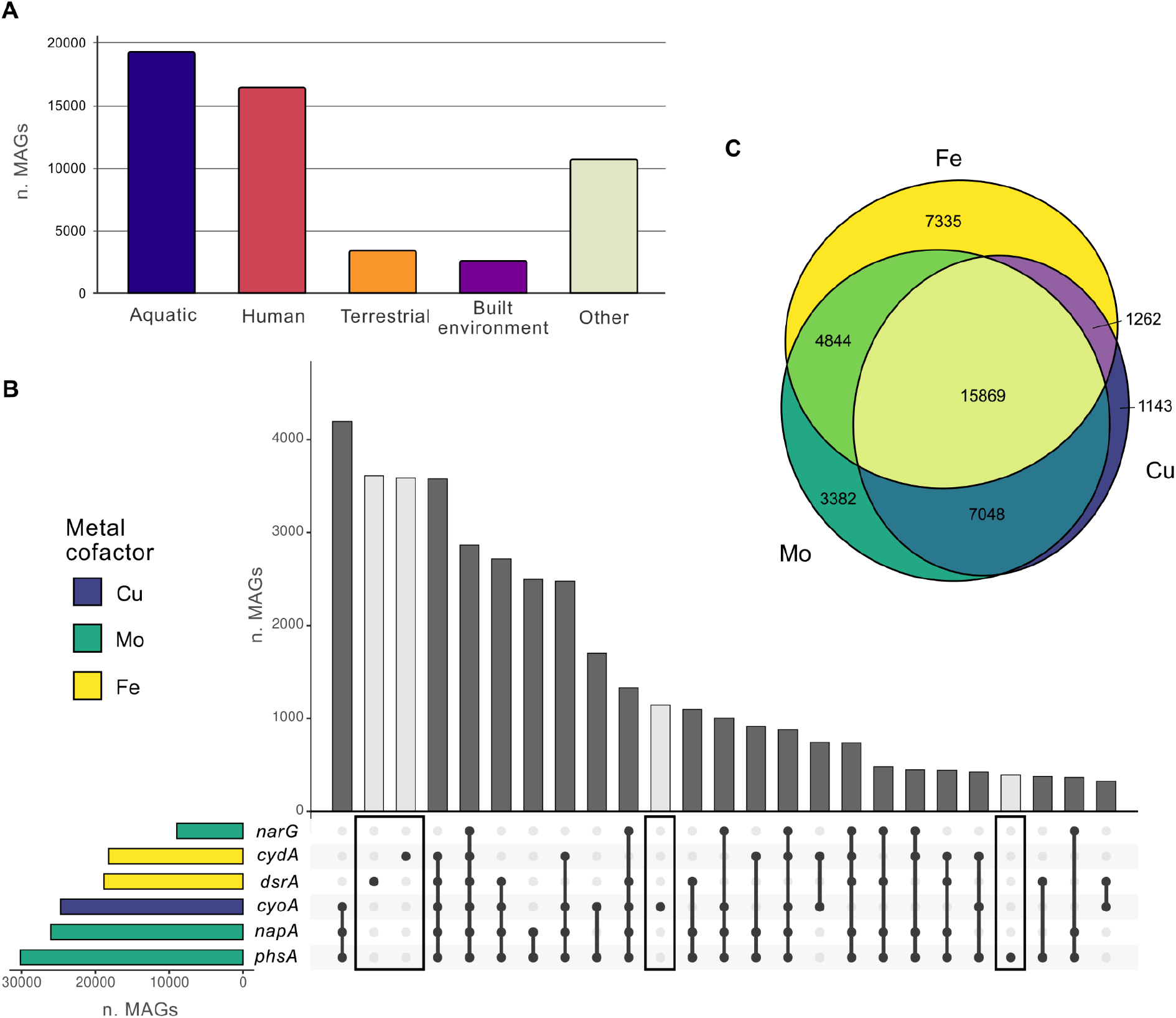
Co-presence of metal dependent oxidoreductases involved in the use of alternative electron acceptors in metagenome assembled genomes (MAGs) annotated from the genomic catalog of Earth’s microbiomes. (*12*). (**A**) Map of the distribution of MAGs analysed for the current study. (**B**) Distribution of oxidoreductases for hydrogen, hydrogen sulfide, ammonia and iron(II) utilisation as electron donor and oxygen, nitrate, nitrite, thiosulfate, elemental sulfur and sulfate utilisation as terminal electron acceptor and their putative metal requirement. (**C**) Venn diagram of the number of MAGs using terminal oxidases requiring copper, iron or molybdenum as cofactors.

In microbiology, the concept of limiting substrate generally refers to a fundamental nutrient that microbial cells consume, and whose availability can limit the growth when it is too low. This definition, originally formulated by Monod, is central to our modern approach to microbiology and microbial ecology (*13, 14*). The most commonly reported examples of limiting substrates include organic and inorganic compounds used as sources of carbon (*13*), oxygen (*15*), nitrogen (*16*), hydrogen (*17*), sulfur (*18*) and phosphorus (*19*), the so-called “macronutrients” that are used as building blocks of the cells. Only recently, the evidence that trace metals or “micronutrients” can also constitute a limiting factor—even when they are not a substrate in chemical reactions—is becoming increasingly clear (*20, 21*). In spite of these observations, the role of trace metals availability in controlling metabolic pathways utilization—independently from substrate availability—is unknown.

Here, we present evidence that trace metals availability can significantly influence growth performance and metabolic shifts in *Escherichia coli* and other microorganisms, including both prokaryotes and eukaryotes. We demonstrate that trace metals deficiencies impose energetic costs on microbial growth, driving shifts toward less energy-dense metabolisms, even when more energetically favourable electron acceptors are present.

### Metabolic Shifts in Response to Trace Metal Availability

*E. coli* can respire oxygen, both in aerobic and microaerophilic conditions, or nitrate under anaerobic conditions (*22*) (Table S3). Under anaerobic conditions with limiting electron acceptor concentrations, *E. coli* performs mixed acid fermentation. These pathways follow a clear thermodynamic hierarchy: oxygen respiration provides the most energy under physiological conditions (ΔG^0^′ = -2,830 kJ/mol glucose), followed by nitrate respiration (ΔG^0^′ = -858 kJ/mol glucose), other possible anaerobic respirations (e. g. fumarate respiration: ΔG^0^′ = -550 kJ/mol glucose), and fermentation (ΔG^0^′ = -218 kJ/mol glucose to acetate) (*22, 23*). The oxidoreductases involved in the utilisation of these terminal electron acceptors require different metal cofactors (Fig. 2A and Table S2). The cytochromes responsible for oxygen reduction require either copper (in the form of a Cu^2+^ ion coordinated by two histidine residues and magnetically coupled to the heme O) for the full aerobic cytochrome bo3 enzyme (CyoABCD) or iron (in the form of several hemes) for the high-affinity cytochrome bd-I and bd-II oxidases (CydAB and AppBC, respectively) used in microaerophilic conditions. Denitrification in *E. coli* involves as many as 5 different enzyme complexes. The first step of denitrification, the reduction of nitrate to nitrite, can be accomplished using either the membrane bound respiratory nitrate reductase (NarGHI and the structurally similar NarZYV) or the high affinity periplasmic nitrate reductase (NapABGHC). Both enzymes, while different in their topology, affinities and operonic organisation, are molybdopterin of the DMSO enzyme class and contain a molybdenum ion coordinated by a two-pterin complex known as molybdopterin (*22, 23*) (Fig. 2A). The subsequent reduction of nitrite to ammonia can be accomplished by two different complexes, either the cytochrome c nitrite reductase (NrfABCD) or the NADH-dependent nitrite reductase (NirBD). Both enzymes rely on iron (in the form of a heme, or a siroheme) as their main catalytic cofactor. During mixed acid fermentation, a suite of diverse oxidoreductases is involved in disposing of reducing equivalents and the release of hydrogen gas, CO_2_ and a variety of low molecular weight organic acids (*24*). These oxidoreductases rely either on iron (for the enzymes alcohol dehydrogenase, AdhE, and succinate dehydrogenase, SdhCDAB), molybdenum (for the formate dehydrogenase of the membrane-bound formate hydrogenlyase (FHL) complex, FdhF, and the nitrate-inducible formate dehydrogenase, FdnGHI), nickel (in the case of the nickel-iron hydrogenase subunit of the formate hydrogenlyase complex, HycE) or magnesium (for the pyruvate dehydrogenase, PoxB) (*25*–*29*) (Fig. 2A). A series of other enzymes, mainly lyases and transferases, that are not involved in redox chemistry, contribute to the formation of the diverse compounds released during fermentation. The relative composition of the compounds released by *E. coli* during fermentation can be altered by changing the primary growth substrate and the growth conditions (*24, 30*).

**Figure 2.**
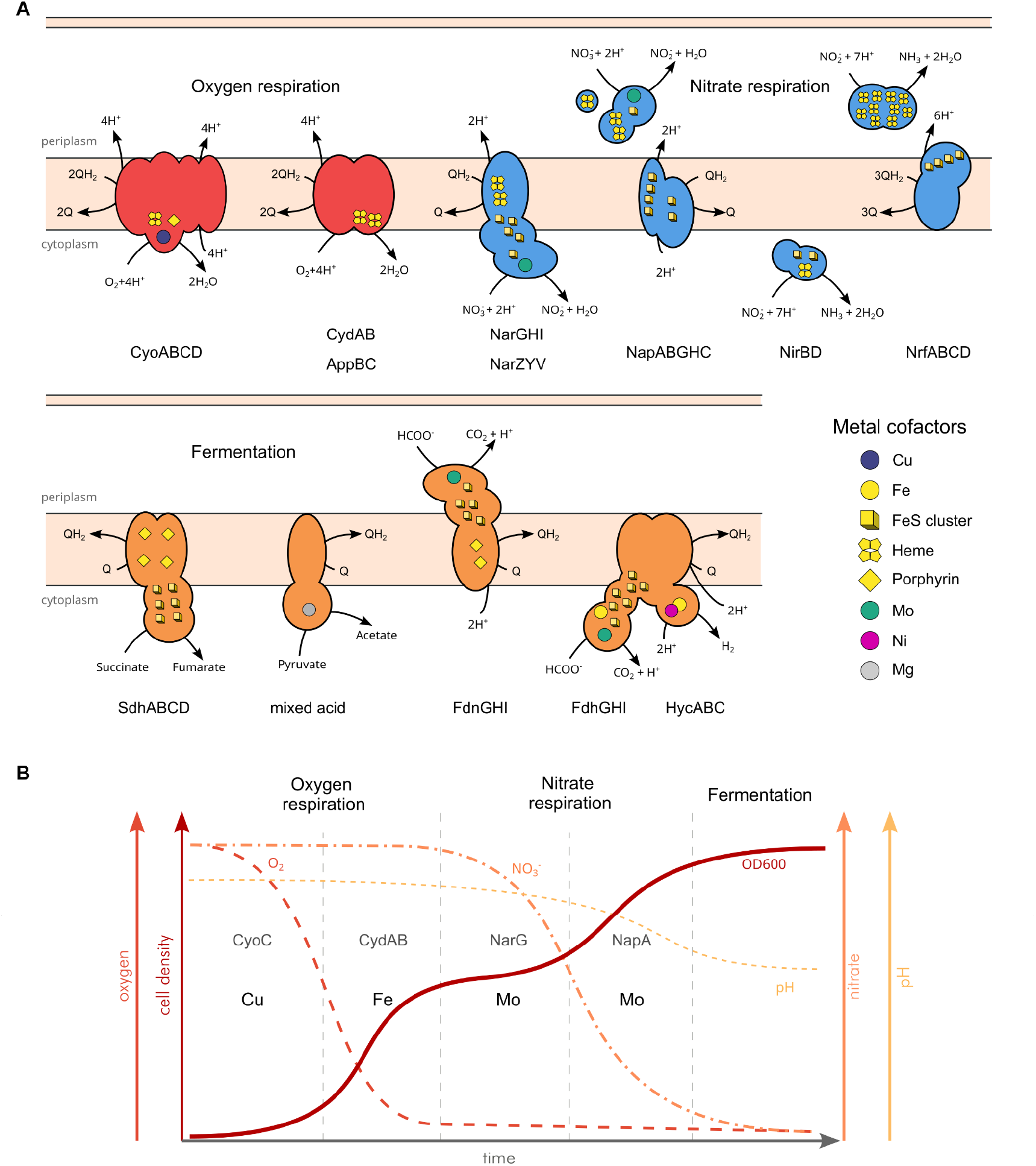
Architecture of *E. coli* metabolic oxidoreductases and expected growth under limiting terminal electron acceptors. (**A**) Topology, architecture and metal cofactors of the oxidoreductases involved in aerobic respiration, denitrification and fermentation. Designation of the enzymes and their subunits is according to the genetic nomenclature and to databases. Underlying data is reported in Supplementary Table 2. (**B**) Expected model of *E. coli* growth under oxygen and nitrate limiting conditions in the presence of the appropriate trace metals. A diauxic growth during the switch from aerobic respiration to denitrification is expected. Altering the availability of Cu, Fe and Mo will change the timing and type of metabolism utilised by the organism.

In minimal media with limiting oxygen and nitrate concentrations, *E. coli* is expected to fully consume oxygen before switching to nitrate reduction, producing ammonia via a nitrite intermediate. Once nitrate is spent, fermentation takes over, producing mixed acids and acidifying the medium (*31*) (Fig. 2B). In the presence of oxygen, the fumarate and nitrate reduction regulatory protein (FNR) is inactivated through oxidation of its FeS cluster, preventing the transcription of the genes for anaerobic respirations (*22*). Alongside with this, oxygen is sensed by the aerobic respiratory control two-component system (ArcBA) which activates and inhibits the transcription of the genes for aerobic and anaerobic respiration, respectively (*32, 33*). As oxygen becomes limited in the environment, the above mentioned mechanisms are inverted and anaerobic respiration genes expression is induced. In the presence of nitrate (and nitrite), NarX and NarQ sensors detect their presence, and activate NarL and NarP, which stimulate transcription of the nitrate (*narGHJI, nap*) and nitrite (*nrf, nir*) reduction genes and repress other anaerobic respiratory systems like fumarate reductase (*frd*) or genes involved in fermentation (*22*). As nitrate levels decrease, fermentation inhibition is released, allowing for its activation. Under this model, oxygen inhibits anaerobic respiration and fermentation, while nitrate presence inhibits fermentation (Fig. S1).

In microbial growth media, trace elements are either added directly as part of the media making recipe, such as in many defined media for chemolithoautotrophic anaerobes, or are provided indirectly as impurities in common salts and from laboratory equipment. We monitored the precise concentrations of trace metals using ICP-MS (see methods for details) in M9 medium (DSMZ 382) made either with common microbiology grade components (BioXtra purity grade) supplemented with a trace metal solution (SL-10, from medium DSMZ 320) or using diverse metal control strategies, including the use of high-purity trace metal grade salts (>99.99 % purity) and ionic exchange high affinity resin (Bio-Rad Chelex 100) for trace metal removal (Table S4 and Fig. S2). As a comparative standard, we tracked *E. coli* metabolic transitions during alternative electron acceptor usage in trace metal (TM)-rich M9 minimal medium with glucose (0.2 % w/v) as sole carbon source by monitoring oxygen, nitrate, pH and cell density (Fig. 3A). Under this condition *E. coli* completely consumed oxygen by 6 hours then switched to denitrification. Nitrate was completely consumed in the media by 12 hours and nitrite accumulation was not visible. Fermentation, as evidenced by the acidification of the medium by ∼1 pH unit, started around 6 hours (Fig. 3A).

**Figure 3.**
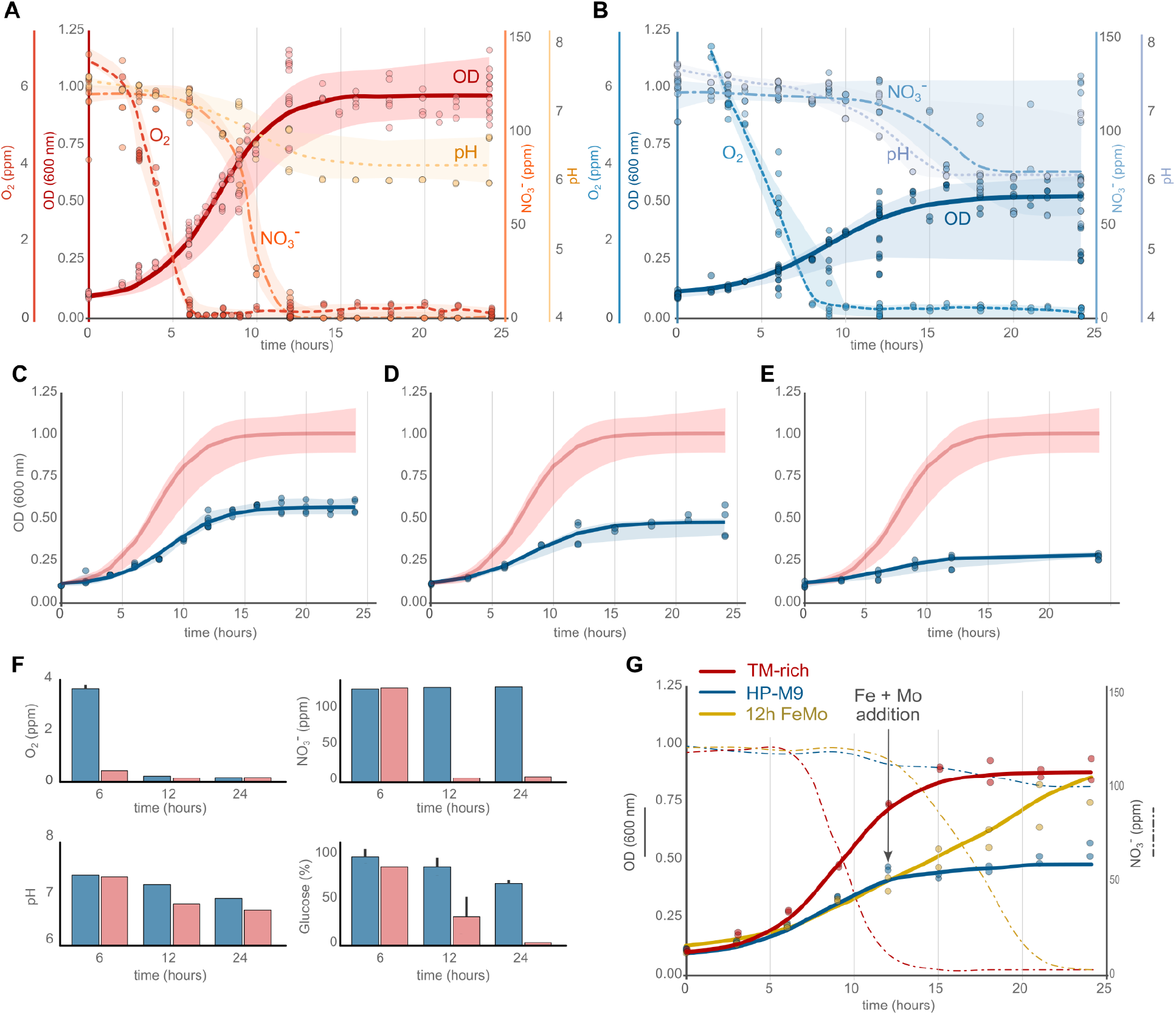
Effect of trace metal availability on the growth and terminal electron acceptor utilization. The lines represent the calculated model (for OD) and averages (for oxygen, nitrate and pH) of all the plotted data while the shaded area represents the area of variability for the data. Individual data points are plotted as circles. (**A**) *E. coli* growth in TM-rich M9 (all experiments), shifting between oxygen, nitrate and fermentation. (**B**) *E. coli* growth in TM-depleted M9 (average of growth in standard M9, high-purity M9 and chelex treated M9 media), switching from oxygen reduction to fermentation while failing to completely consume nitrate. (**C, D** and **E**) *E. coli* growth in anaerobiosis in response to varying trace metal availability (standard M9, high-purity M9 and chelex treated M9, respectively). (**F**) Oxygen, nitrate, pH and residual glucose in the media for *E. coli* grown in TM-rich and TM-depleted (chelex treated M9) conditions at 6, 12 and 24 hours. Each data point represents the mean ± the standard deviation of three independent samples. (**G**) The addition of molybdenum and iron at 12 hours during growth restores nitrate consumption and growth to TM-rich levels.

In trace metal (TM)-depleted media (Table S4), *E. coli* exhibited slower growth, reaching only ∼45 % in high-purity M9 (HP-M9) and ∼25 % in chelex-treated M9 (chlx-M9) of the cell density observed in TM-rich conditions (Fig. 3B-D). The absence of trace metals not only reduced overall growth but also affected the timing of electron acceptor utilisation. Indeed, oxygen consumption was delayed by 3 hours in HP-M9 and 6 hours in chlx-M9 compared to TM-rich medium. Nitrate reduction never began in TM-depleted chlx-M9 medium (Fig. S3), whereas it started between 9 and 12 hours in HP-M9, and nitrate was never fully consumed, remaining stable at about 92 % of the initial concentrations throughout the entire experiment (ended at 24 hours). In both TM-depleted media fermentation-induced acidification began while nitrate was still present, suggesting that trace metal deficiency affects the metabolic switch in *E. coli* beyond substrate availability. To assess whether the incomplete utilization of nitrate was the result of specific trace metal depletion, we tested double metal complementation at 12 hours to rescue nitrate respiration. The addition of molybdenum and iron at 12 hours fully restored nitrate reduction (Fig. 3G). Similar results were obtained in fully aerobic cultures grown in M9 TM-rich and TM-depleted media with a growth rescue obtained by complementing with the addition of copper and iron (see Fig. S5 and Supplementary text). These results demonstrate that the availability of specific trace metals used by oxidoreductases as key metal cofactors control metabolic shifts in the utilization of alternative electron acceptors.

In *E. coli*, molybdenum availability holds a primary role in the transition from aerobic to anaerobic respiration (*34*). Many key enzymes for anaerobic respiration, including the different nitrate reductase, the trimethylamine N-oxide (TMAO) reductase and the dimethylsulfoxide (DMSO) reductase, are molybdoenzymes—enzymes that incorporate a molybdenum ion in their active site within a molybdenum pterin cofactor (Moco). During anaerobiosis, Moco biosynthesis in *E. coli* is activated by the transcription factor FNR, which induces the expression of the *moaABCDE* operon, essential for the production of active molybdoenzymes (*35*) (Fig. S1). This process is iron-dependent since both FNR and the MoaA protein require iron-sulfur clusters for their activity (*36*). Furthermore, molybdenum plays a direct role in regulating the expression of anaerobic respiration pathways; for example, nitrate respiration genes are activated by the NarX-NarL two-component system, which responds to the presence of both nitrate and molybdenum ion (*37*). These regulatory mechanisms, when considered in the light of our results, provide direct evidence that the availability of suitable trace metals controls shifts in electron acceptor utilization. In the absence of molybdenum, only nitrite and fumarate respiration remains feasible together with fermentation, since the key enzymes for these pathways require only iron, forcing a fast shift towards less energy-efficient pathways.

To understand the cellular mechanisms underpinning these responses to trace metal scarcity, we combined physiological measurements with label-free proteomics and functional assays. Growth yields (calculated as the increase in biomass per gram of glucose consumed) show that in the TM-depleted conditions *E. coli* converts only 6.0 % and 5.4 % of the glucose to biomass, during anaerobic and aerobic growth respectively, compared to 23.3 % and 36.5 % in TM-rich conditions. A significant portion of the consumed glucose is probably used for cell maintenance and activities other than cell growth in TM-depleted conditions, as suggested by the percentage of glucose consumed not converted to biomass that is 76.7 % and 63.5 %, for anaerobic and aerobic growth, respectively (Fig. 3 and S4), compared to ∼30 % for the TM-rich condition. We hypothesised that in the absence of trace metals, *E. coli* would preemptively express oxidoreductases for alternative electron acceptors while upregulating metal transporters to compensate for the metal limitation. This upregulation of the metal acquisition machinery could account for the additional energy not converted to biomass.

Proteomic analysis of TM-rich and TM-depleted anaerobic cultures at 6, 12, and 24 hours (n =18) supported this hypothesis (Fig. 4 and Fig. S6 and S7). In TM-depleted cultures, enzymes involved in nitrate reduction displayed a 2 to 30-fold decrease (Fig. 4C and 4D), while some of them, like the membrane bound nitrate reductase NarG and the periplasmic nitrate reductase NapA, were completely undetectable at all time points. Fermentation enzymes, despite showing a quite stable expression in both conditions, were less affected by TM depletion. Additionally, many metal uptake and homeostasis proteins could be exclusively detected in samples from TM-depleted cultures (Table S5), and the differential analysis highlighted a 2 to 500-fold increase in their expression compared to TM-rich cultures (Fig. 4D and 4E, Table S5). Siderophore production assays further confirmed enhanced metal scavenging under TM-depleted conditions, with siderophores levels being undetected in the TM-rich condition while consistently increasing over time in the TM-depleted cultures (Fig. S8). The temporal evolution of the proteome in the two conditions shows a rapid metabolic switch in *E. coli* grown in TM-depleted condition and the upregulation of the metal acquisition machinery. The trace metal induced metabolic shift in the TM-depleted culture is evident when investigated differentially expressed protein clusters (see Fig. S9 - S11 and Supplementary Text). These results show that trace metal depletion, besides controlling electron acceptor utilization, imposes a large energetic cost on the cell.

**Figure 4.**
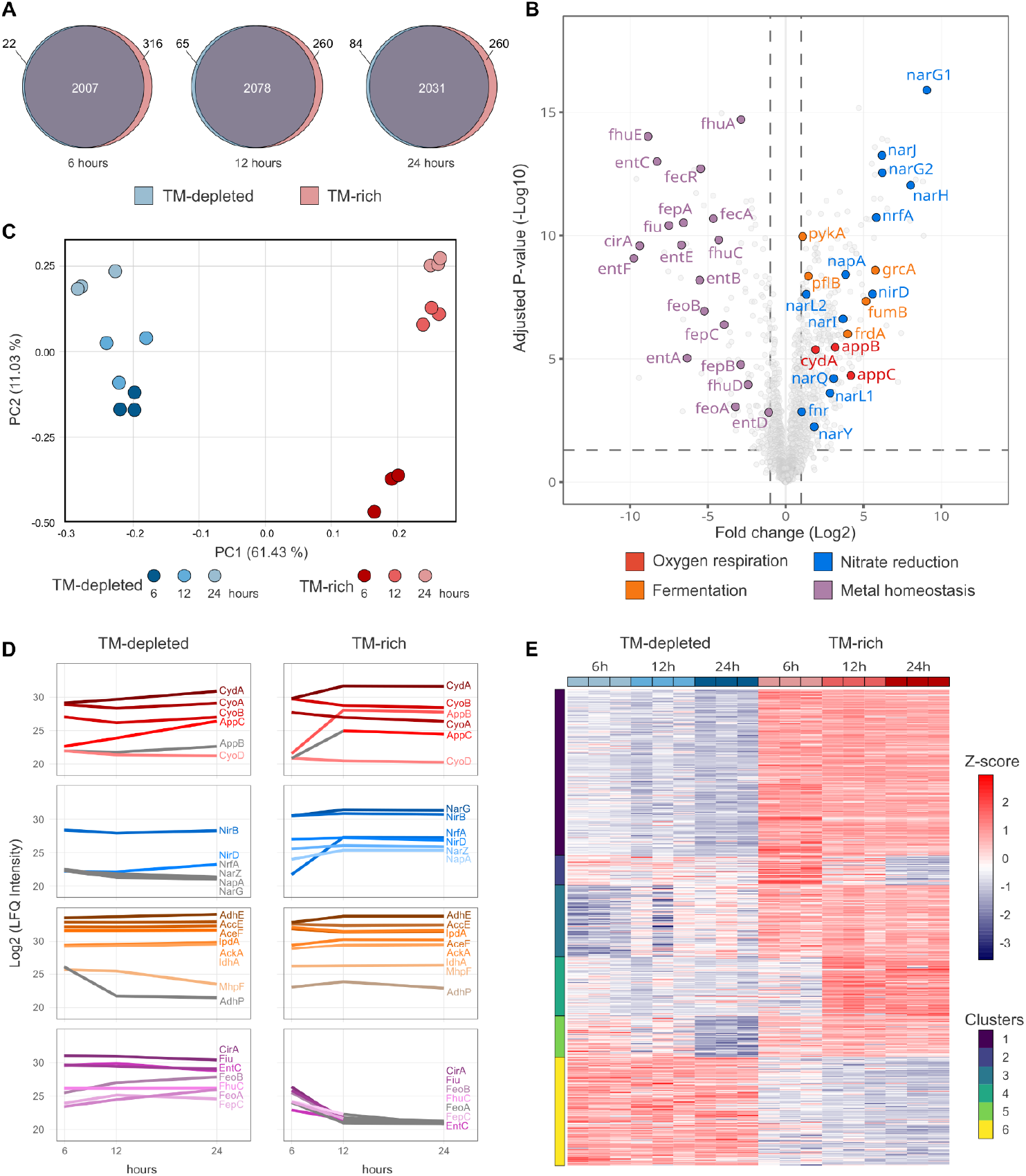
Proteomic analysis of the *E. coli* grown in TM-depleted and TM-rich conditions. (**A**) Venn diagram representing shared and unique proteins between the two conditions at 6, 12 and 24 hours. (**B**) Volcano plot of the differentially expressed proteins at 12 hours between the two growth conditions. (**C**) Principal component analysis of the differentially expressed proteins for each of the biological replicates. (**D**) Temporal variations of the LFQ intensities for the proteins involved in oxygen respiration (top row), nitrate respiration pathway (second row), fermentation (third row) and metal homeostasis (bottom row) between the two growth conditions at 6, 12 and 24 hours. Segments in gray indicate that the proteins could not be detected in any of the biological replicates for a given condition and imputed values were used in the construction of the plot. (**E**) Heatmap of the differentially expressed proteins (Z normalized) across all samples.

To determine whether similar mechanisms occur in other microorganisms, we tested *Bacillus subtilis* (a Gram-positive bacterium), and *Stichococcus bacillaris* (a green alga) under TM-rich and TM-depleted conditions (Fig. S12 and S13). Both organisms exhibited reduced growth in TM-depleted media. *B. subtilis* displayed strong effects during aerobic growth (Fig. S12A) and a similar delayed switch between electron acceptors, with nitrate reduction limited by molybdenum and iron deficiencies (Fig. S12B). The green alga *S. bacillaris* displayed reduced biomass (∼51 % compared to TM-rich media, Fig. S13A) and evidence of photosystem decoupling under TM-depletion (Fig. S13B), suggesting inefficient energy utilization and accumulation of reducing equivalents in the photosystem. In *B. subtilis*, the substrate-to-biomass conversion rates were lower in TM-depleted conditions, indicating increased energy costs due to the scarcity of necessary trace metals, similar to what we observed in *E. coli* (Fig. S12C-D).

### Ecological and Evolutionary Implications

Thermodynamic principles predict the order in which diverse electron acceptors are utilised by microbes in natural systems. In marine sediments, for example, oxygen is used first, followed by nitrate, nitrite, manganese oxide, ferric iron, and sulfate until only CO_2_—a weak electron acceptor—remains (*38*). While thermodynamics helps predict the most high-energy redox couples, it does not provide a complete picture, as it cannot account for important factors like reaction kinetics, the metabolic costs of the required molecular machinery and the availability of the necessary cofactors to drive these reactions.

The availability of metal ions represents a crucial factor for the health, functions, and dynamics of ecosystems (*4*). Our findings suggest that trace metals may influence microbial competition by controlling access to specific redox couples, regardless of substrate availability. This might provide a possible explanatory mechanism for the high functional redundancy encountered in natural ecosystems (*39, 40*), and drive competition for resources beyond the classic models connected with substrate availability and specificity (*4*). Recent data suggest how cofactor utilization can shape microbial communities. For example, the differential Fe and Cu availability has been proven to shape phytoplankton metabolism and primary productivity in the surface ocean (*41*). Similarly, the partitioning of ammonia oxidizing bacteria and archaea in the oceanic water column has been linked to their differential utilization and capability to acquire iron and copper, respectively used in the key enzyme ammonia monooxygenase (*42*).

Our survey of metagenome assembled genomes from diverse ecosystems (Table S1) suggests that the ability to use alternative electron acceptors is widespread in taxonomically diverse microorganisms (Fig. 1B). These mechanisms play a key role in the survival and structuring of marine microbial communities, allowing organisms to respond to fluctuating availability of electron acceptors (and donors). Additionally, our data show that up to 70.6 % of the investigated genomes encodes for more than one terminal reductase requiring different trace metals as cofactors (Fig. 1B). Specifically, ∼37.9 % of the genomes encode for reductases that require iron, copper or molybdenum (Fig. 1C). Given that trace metals are not uniformly distributed in the environment, this data shows that the ability to use diverse metal cofactor might provide a competitive advantage and might be a key factor structuring ecological niches.

Despite this, the availability of suitable substrates has long been considered the key factor controlling competition and niche partitioning (*43, 44*). Our data show that the availability of metals used as cofactors imposes strict controls on redox couple utilization, independent from substrate availability (Fig. 3). A complex regulatory network links substrate availability to metal cofactor availability in *E. coli* (Fiig. S1), and similar mechanisms might be present in most microorganisms (*23, 45*). Based on our results, we speculate that microorganisms with greater plasticity in their use of cofactors and more sophisticated metal uptake mechanisms might have an advantage under trace metal limiting conditions also when competing for the same substrate. This is well established in opportunistic pathogens, like *Pseudomonas aeruginosa*, where siderophores are considered virulence factors, affording the bacterium a competitive edge in the colonization of the Fe-depleted host (human) tissues (*46*).

Our results offer direct evidence of how trace metal availability might have influenced the evolution of metabolism and biogeochemistry. Changes in trace metal availability over geological time, such as those following the Great Oxidation Event (GOE) and Neoproterozoic Oxygenation Event (NOE) (*47*), have likely shaped the evolution of microbial metabolisms. Comparative structural protein analysis have suggested that the diversity of metal binding folds has evolved very early during life evolution (*3, 11, 48, 49*), suggesting that fold innovation happened early and that metals were acquired during evolution by substitution within existing protein folds (*48*). Given that early life has been suggested to lack both the structures required to control intracellular metal concentrations (*48*), the utilization of trace metal as cofactor was strongly controlled by their environmental availability. Our results provide direct evidence of how trace metal availability affects metabolic shifts and the energetic costs connected with their scarcity. It is possible that these mechanisms might have provided selective pressure to drive metabolic innovation and evolution, driving the evolution of biogeochemistry (*4*). The environmental distribution of metals has changed as the result of geological processes acting on diverse temporal and spatial scales (*50*). Complex feedbacks between geological mechanisms providing trace metals and the evolution of more sophisticated strategies to acquire and use available metals by life likely pushed diversification and geosphere-biosphere coevolution.

Taken together, our results demonstrate that trace metal deficiency imposes strong energetic costs on microbial growth, driving the shift toward less energy-dense metabolisms even when more energetically favourable electron acceptors are present. Understanding how trace metals influence microbial functional diversity in the environment may unlock new strategies for managing microbial communities in environmental and industrial settings.

## Funding

This work was supported by funding from the European Research Council (ERC) under the European Union’s Horizon 2020 research and innovation program Grant Agreement No. 948972, acronym COEVOLVE to DG. BdP was supported by funding from European Union’s Horizon Europe research and innovation programme under the Marie Skłodowska-Curie grant agreement No. 101154017, acronym Subcarb.

## Author Contributions

Conceptualization: DG, CV, AR, BdP

Methodology: DG, AR, BdP, MaCa, AP, NN, MoCo, CV, SL, SC, RoIa, RaIs, MM, JB

Investigation: AR, BdP, MG, MaCa, NN, MoCo, LdI, SC, RoIa, SL, JB

Visualisation: DG, AR, BdP, SL

Funding acquisition: DG

Project administration: DG, AC

Supervision: DG, AC, AP, CV, SL, RaIs, MM

Writing – original draft: DG, AR, BdP

Writing – review & editing: All authors

## Competing interests

The authors declare no competing interest.

## Data and code availability

All the data and the codes supporting this work is open source and can be found in the following link: https://github.com/giovannellilab/Ecoli_trace_metal_metabolic_shift and has been released in Zenodo under the DOI 10.5281/zenodo.14563099. The Escherichia coli genome is available through ENA bioproject PRJEB85325. The proteomic data has been deposited in the PRoteomics IDEntifications (PRIDE) database at the European Bioinformatics Institute with accession number PXD059598. The MAGs are publicly available and were downloaded from the JGI GEMs Genomic catalogue of Earth’s microbiomes (https://genome.jgi.doe.gov/portal/GEMs/GEMs.home.html).

## Supplementary Materials

Materials and Methods

Supplementary Text

Figs. S1 to S12

Tables S1 to S10 References (51–74)

## Notes

https://github.com/giovannellilab/Ecoli_trace_metal_metabolic_shift

https://doi.org/10.5281/zenodo.14563099

https://www.ebi.ac.uk/ena/browser/view/PRJEB85325

https://www.ebi.ac.uk/pride/archive/projects/PXD059598

### Competing Interest Statement

The authors have declared no competing interest.

### Summary of Updates

Improved the supplementary figures and minor changes to the main manuscript to improve readability

