## Supplementary Online Materials for "Trace metals availability controls terminal electron acceptor utilization in *Escherichia coli*"

**The PDF file includes:**

Materials and Methods

Supplementary Text

Figs. S1 to S12

Tables S1 to S10

References (51–74)

#### Materials and Methods

##### Culture media and growth conditions

*Escherichia coli* K12 DH5α (*E. coli*) from our collection was used as a model organism for the physiological studies carried out in this work. *E. coli* was grown at 37 °C shaking at 180 rpm in TY medium (DSMZ medium 1143, Tables S6) for the preparation of the pre-innocula, at 37 °C shaking at 180 rpm in M9 medium (DSMZ medium 382, Tables S7) for aerobic and anaerobic growths.

To test different levels of metal contaminations, four different M9 media were prepared: 1. M9 medium as defined by DSMZ (M9) using non metal free labware, BioXtra purity grade salts and deionized water (high level of trace metal contaminations, but no added trace metals); 2. M9 medium high purity (HP-M9) using acid-washed labware, Merck group trace metal basis grade salts (>99.99 % purity) and ultra-pure water; 3. M9 medium high purity subjected to a chelation ion chromatography (chlx-M9) to remove all metal impurities. Metal chelation was achieved using ion exchange resin (see below for details) 4. M9 medium supplemented with trace element solution SL-10 (from DSMZ medium 320, Tables S8) prepared with trace metal basis grade salts (>99.99 % purity) (TM-rich).

For all *E. coli* growth experiments, an overnight culture in TY medium was diluted to an optical density of 0.1 OD (at 600 nm) in the selected M9 medium, and grown in aerobic condition for 24 h to pre-adapt *E. coli* to the different media and make it more sensitive towards trace metal availability changes. The pre-adapted cultures were then centrifuged and resuspended in different volumes of fresh media to an $OD_{600}$ of 0.1 and inoculated in 250 ml glass flasks, with a final volume of 50 ml of culture, for aerobic growth experiments, or in 120 ml crimp-top glass vials, with a final volume of 80 ml of culture, for anaerobic growth experiments. In both experiments, cell growth was monitored for 24 h by measuring optical density at six selected time points (0, 3, 6, 9, 12, and 24 h). At these same time points, an aliquot of 1 ml from each culture was centrifuged and filtered through a 0.22 µm cutoff filter to remove the cells, and the resulting cell-free supernatants were analysed to measure pH, glucose, nitrate and dissolved oxygen concentrations, as described below.

*Bacillus subtilis* PY79 (*B. subtilis*) from our collection was used as a model Gram-positive organism in this study. To prepare the pre-inocula of aerobic growth, *B. subtilis* was grown in TY medium (DSMZ medium 1143, Tables S6) at 37 °C overnight with shaking at 180 rpm. An overnight culture in TY medium was diluted to an optical density of 0.1 OD (at 600 nm) in the selected M9 or TE medium and grown in aerobic conditions for 24 h to pre-adapt *B. subtilis* to the different media and make it more sensitive towards trace metal availability changes. The pre-adapted cultures were centrifuged and resuspended in fresh media to an $OD_{600}$ of 0.1 and inoculated in 500 ml glass flasks. Cell growth was monitored for 24 h by measuring optical density at six selected time points (0, 3, 6, 9, 12, 24, and 30 h). At these same time points, an aliquot of 1 ml from each culture was centrifuged and filtered through a 0.22 µm cutoff filter to remove the cells, and the resulting cell-free supernatants were analyzed to measure pH and glucose.

The model eukaryotic organism *Stichococcus bacillaris* Nägeli (Trebouxiophyceae, Chlorophyta), strain ACUF 102, obtained from the Algal Collection at University of Naples Federico II (http://www.acuf.net/), was used in this study. A pre-adapted culture of *S. bacillaris* was grown in Bold's Basal Medium (BBM) (Table S9 and S10) at 24 °C, under continuous shaking at 120 rpm. Illumination was provided at an intensity of 80 µmol photons $m^{-2}$ $s^{-1}$. To analyse the effect of trace metals availability on the eukaryotic model organism, 5 ml of pre-adapted culture of *S. bacillaris* in the exponential phase were resuspended in fresh BBM media supplemented (TM) or depleted (BBM) of trace metals. Growth was monitored daily by spectrophotometric measurements at a wavelength of 550 nm. A calibration curve was established to determine the linear coefficient between optical density and mass concentration.

All growth experiments were performed in 3 biological replicates.

**Acid washing and metal removal by ion exchange chromatography**

In order to control trace metal contamination, all laboratory equipment, plasticware and glassware were acid washed prior to use. Briefly, all material was washed in laboratory CITRANOX acid detergent (Sigma) thoroughly before performing several rinses in trace-metal free deionized water. All material was then immersed in a fresh solution of 1 % $HNO_3$ made from high purity acid (ROMIL-SpA™ Super Purity Nitric Acid 67-69 %) for at least 24 hours. The material was then washed three times using metal-free Milli-Q water (18 Mohm) and air dried in the chemical hood of our geochemical and trace metal laboratory. We routinely run blanks in the lab using water and acid solution (see metal concentration determination section).

The removal of additional metal ions was achieved through ion exchange chromatography. Briefly, 5 grams of Chelex 100 resin (Bio-Rad, California, United States) were resuspended in ultra-pure water and poured into a 50 ml glass chromatography column. Several bed volumes of ultrapure metal-free Milli-Q water were run through the column before running the HP-M9 media to ensure proper packing. A solution of M9 salts 10-fold concentrated was gradually flown through the column and collected in acid-washed glass bottles. Run was performed at room temperature and by gravity flow with the running sample positioned 50 cm above the column. Same protocol has been performed on M9 medium components $CaCl_2$ (100 mM), $MgSO_4$ (1 M) and glucose (40 %).

**Dissolved oxygen, nitrate, pH and glucose determination**

Dissolved oxygen was monitored through a non-invasive measurement procedure using oxygen sensor spots SP-PSt3-NAU and a stand-alone fiber optic oxygen meter Fibox 4 (PreSens - Precision Sensing GmbH, Regensburg, Germany). The oxygen spots were attached to the inner surface of 120 ml crimp-top glass vials used for anaerobic cultures by using a transparent silicone glue. During growth, dissolved oxygen was measured at selected time points in three technical replicates for each biological replicate and reported as a concentration in parts per million (ppm).

Nitrate consumption was monitored during growth by ion chromatography by using a ECO IC (Metrohm, Switzerland) equipped with a Metrosep A Supp 5 column (Metrohm) for anion separation. Determination was carried out on cell-free supernatants obtained by centrifugating the cultures for 5 min at 8,000 × g and 4 °C (HERMLE Labortechnik Centrifuge Z216MK, Germany, equipped with a 220.87 rotor). Three technical replicate measurements were carried out for each biological replicate. Sample preparation and analytical procedures have been performed as reported in the Standard Operating Procedure (*51*).

pH variations of the culture media during growth were monitored using the bromothymol blue (BTB) spectrophotometric assay (*52*). BTB is a commonly used acid-base indicator, with pKa ≈ 7.5. At pH values above 7, BTB is stable in a blue form with an absorption maximum at 620 nm, whereas at reducing pH, BTB switches to a protonated yellow form, which absorbs at 430 nm. pH measurements of *E. coli* supernatants were made by mixing 75 µl of each sample with 25 µl of BTB solution at a final concentration of 0.01 % in the wells of a 96-well plate. The decrease of the blue color at reducing pH was measured by reading the absorbance at 620 nm. A calibration curve in the range of pH 6 to 8 was run in parallel to infer the samples pH values from measured absorbance values. Three technical replicate measurements were carried out for each biological replicate. Selected points were verified using a laboratory pH-meter regularly calibrated.

The concentration of glucose was measured using the Megazyme D-Glucose GOPOD kit (Megazyme) according to manufacturing procedure. Briefly, the diluted samples properly prepared for analysis (glucose range between 0.04-1 g/L) were mixed with GOPOD solution. The solution mixtures were incubated for 20 min at 45 °C. Absorbance was measured using a Cary 3500 Multicell UV-Vis Spectrophotometer (Agilent) at 510 nm. Each sample was prepared in duplicate.

**PAM Fluorometry Assay**

The photosynthetic efficiency of the *S. bacillaris* culture was assessed at 14 days using a FMS2 Hansatech Pam fluorometer. Samples were adapted in the dark for 30 minutes and then transferred into a 4 ml quartz glass cuvette containing a magnetic micro-agitator to ensure homogeneity during tests (White et al., 2011). Saturating light was applied to measure F0 (Minimum fluorescence yield) and Fm (Maximum fluorescence yield), thus maximum quantum efficiency of photosystem II was calculated according to the following equation:

$$Fv/Fm= (Fm-F0)/Fm$$

A sequence of 9 increasing actinic irradiance, randing from 36.56 μEin to 1213 μEin was set in order to evaluate Fs (Fluorescence yield in the steady-state in light-adapted samples), F’0 ( Minimum fluorescence yield in light-adapted samples), F’m (Maximum fluorescence yield in light-adapted samples). Before each actinic irradiance, the samples were incubated at dark for 2 minutes.

**Growth curve and data analysis**

Growth curves data were analysed in R (*53*) using the Growthcurver package (*54*). Briefly, a logistic model was fitted to each experimental run, consisting of at least three biological replicates, and multiple experimental runs were also pooled together to get a model reflecting the variability obtained during the entire experimental work. Model parameter growth rate, doubling time, and the area under the logistic curve were used to compare the different experiments. The obtained figures were combined with data from oxygen and nitrate concentrations and pH in inkscape (https://inkscape.org/) to obtain the final figures.

**Genome sequencing**

The genome of the *E. coli* strain K12 DH5α used for the experiments was sequenced to obtain a high quality reference genome for the proteomic analysis. The genome was sequenced by MicrobesNG (Birmingham, United Kingdom) using a combination of Illumina and Oxford Nanopore Technologies (ONT) sequencing. High quality genomic DNA from a fresh culture of E. coli grown in TY medium was obtained using a modified phenol:chloroform extraction (*55*), visualized on agarose gel and shipped for sequencing. Before library preparation, DNA was quantified using the Quant-iT dsDNA HS (ThermoFisher Scientific) assay in an Eppendorf AF2200 plate reader (Eppendorf UK Ltd, United Kingdom) and diluted as appropriate. Genomic DNA libraries were prepared using the Nextera XT Library Prep Kit (Illumina, San Diego, USA) following the manufacturer’s protocol with the following modifications: input DNA was increased 2-fold, and PCR elongation time was increased to 45 seconds. DNA quantification and library preparation were carried out on a Hamilton Microlab STAR automated liquid handling system (Hamilton Bonaduz AG, Switzerland). Libraries were sequenced on an lllumina NovaSeq 6000 (Illumina, San Diego, USA) using a 250 bp paired end protocol. Reads were adapter trimmed using Trimmomatic version 0.30 (*56*) with a sliding window quality cutoff of Q15. To improve genome assembly, DNA libraries were also prepared with Oxford Nanopore Technologies (ONT) SQK-RBK114.96 kit (ONT, United Kingdom) using 200-400 ng of High Molecular weight (HMW)

DNA. The sequencing library was loaded in a FLO-MIN114 (R.10.4.1) flow cell in a GridION (ONT, United Kingdom). Hybrid assembly of ONT and Illumina reads was performed using Unicycler version 0.4.0 (*57*), and contigs were annotated using Prokka 1.11 (*58*).

#### Metagenome Assembled Genomes annotation

Metagnome Assembled Genomes (MAGs) were downloaded from the JGI GEM database (https://genome.jgi.doe.gov/portal/GEMs/GEMs.home.html) and consisted of 52,515 metagenome-assembled genomes from over 10,450 metagenomes collected from diverse microbiomes (*12*). The MAGs are of medium-quality level in the MIMAG standard (mean completeness = 83 %, mean contamination = 1.3 %). MAGs were annotated using Hmmer3 (*59*), by searching a series of HMM models obtained from InterPro (*60*) coding key terminal reductases (Table S2). Custom python and R scripts together with the rhmmer package (https://github.com/arendsee/rhmmer) were used to summarize the results. The code and data tables are available in the GitHub repository.

#### Siderophore production measurement

Siderophore production was assessed using Chrome Azurol S (CAS), and following a quantitative procedure (*61*, *62*). The CAS solution was prepared as follows: 1.5 ml of a 1 mM $FeCl_3$ solution in 10 mM HCl was mixed with 7.5 ml of a 2 mM CAS solution and slowly added to 46 ml of a 1.5 mM CTAB solution with constant stirring. To this mix, 32 ml of a piperazine acid solution (1.6 M anhydrous piperazine in 2.4 M HCl) was added gradually, and the solution was adjusted to a final volume of 100 ml. Finally, 87.3 mg of 5-sulfosalicylic acid was dissolved into the solution, to achieve a final concentration of 4 mM. For the assay, 100 µl of culture supernatants were mixed with 100 µl of the prepared CAS solution in a 96-well microplate and incubated in the dark for 1 hour. Absorbance was measured at 620 nm. Controls included 100 µl of CAS solution mixed with EDTA 200 mM (positive control), Milli-Q water, and uncultured growth media (negative controls). Siderophore production is expressed as a siderophore production unit (psu) normalized to the positive control (EDTA), using the following equation:

$$psu = [(A_{620,sample} - A_{620,media}) / (A_{620,EDTA} - A_{620,milliQ})] \cdot 100$$

#### Metals quantification

Trace metal concentrations in the different media were analyzed by Inductively Coupled Plasma Mass Spectrometry (ICP-MS) using an ICP-MS 7900 (Agilent, California, United States). Sample preparation and analytical procedures have been performed as reported in the Standard Operating Procedure (*63*).

Briefly, all samples were filtered using 0.22 µm cutoff filter and diluted in 1 % $HNO_3$, which also serves as blank. Operational parameters, including argon flow rate, radiofrequency power, and ion lens configuration, were optimized to ensure precision. The prepared samples were then analysed with the ICP-MS instrument. Calibration curves were carried out using certified standards. Data acquisition and analysis were conducted using ICP-MS MassHunter 4.6 software version. A series of blanks and standards was routinely run during the experiments to verify the absence of element carryover and to check for peak drift. These include two quality control blanks analysed as samples at the beginning of the run, along with one blank and one standard every 10 samples analysed.

**Cell lysis, sample preparation and proteomic**

For proteomic sample preparation, cells grown in TM-depleted and TM-rich conditions were centrifuged at 2500 × g and 4 °C for 10 min (Neya 16R, REMI, India, equipped with a A 6-50 rotor) at different time points (6, 12, and 24 h) and frozen at -80 °C until lysis. Three biological replicates per time point were used, for a total of 18 different proteomes. The cells were resuspended in 1 ml lysis buffer (50 mM Tris HCl pH 8, 150 mM NaCl, 6 M urea, 2 % SDS, 10 mM EDTA, 10 mM DTT) with added protease and phosphatase inhibitors (Protease Inhibitor Mix M, SERVA electrophoresis Gmbh, Heidelberg, Germany), and frozen and thawed three times and sonicated on ice at a setting of 40 % amplitude for 10 min (30 sec on, 30 sec off) (Vibra cell VCX 130, Sonics, USA, equipped with a 6 mm probe). Crude extracts were then centrifuged for 30 min at 6000 × g, 4 °C to remove the insoluble fraction. Protein concentrations were measured using a Pierce™ BCA Protein Assay Kit (Thermo Fisher Scientific) following the manufacturer's instructions. The calibration curve was performed preparing the standard (Bovin Serum Albumin, BSA) in the lysis buffer to avoid interferences. Protein samples were subjected to label-free LC-MS/MS shotgun proteomics (Functional and Structural Proteomics Facility, CEINGE–Biotecnologie Avanzate, Italy). Briefly, protein extracts were first mixed with SDS (5 % final concentration) and DTT (20 mm final concentration), boiled, cooled to room temperature, and then alkylated with iodoacetamide (40 mm final concentration) in the dark for 30 min. Afterward, phosphoric acid (at a final concentration of 1.2 %) and six volumes of binding buffer (100 mM triethylammonium bicarbonate (TEAB) in 90 % methanol) were added to each sample. After gentle mixing, the protein solutions were loaded to an S-Trap filter, spun at 2000 rpm, and the flow-through collected and reloaded onto the filter for three times, and then each filter was washed with binding buffer three times. Finally, a digestion buffer containing trypsin at 1:10 wt:wt in 50 mM TEAB was added into the filter and the samples were incubated o/n at 37 °C. Peptides were eluted according to the manufacturer instructions and the peptides-containing fractions were pooled, lyophilized and resuspended in 0.2 % Formic Acid. LC-MS/MS analyses were performed on a Vanquish-Neo UHPLC system coupled to an Orbitrap Exploris 240 (ThermoScientific, Waltham, MA).

After loading, the samples were first concentrated and desalted on the pre-column (PepMap™ Neo Trap cartridge) and then fractionated on a C18 reverse phase capillary column (Double nanoViper™ PepMap™ Neo 2 µm C18, 75 µm X 150 mm) using a gradient from 5 % to 95 % of solvent B (0.2 % formic acid in 95 % acetonitrile) in A (0.2 % formic acid and 2 % acetonitrile in ultrapure water) over 77 min, at a flow rate of 250 nl $min^{-1}$. The MS/MS acquisition was performed in data dependent mode (DDA) mode by fragmenting the 10 most intense ions in the collision-induced dissociation (CID) modality with a full scan in the 300 to 1800 m $z^{-1}$ range. Three technical replicates were run for each sample.

**Differential proteomic analysis**

For proteomics data processing, raw files were converted to the open mzML format with the msConvert tool of ProteoWizard (*64*) and then subjected to an optimized label-free quantification pipeline (*65*). In detail, DDA data were analyzed with MSFragger (v4.1)(*66*) in FragPipe (v22.0) with the default LFQ workflow with match between runs, using as reference database for protein identification the newly sequenced genome of *E. coli* K12 DH5α (NCBI bioproject PRJNA1129478). A quantification matrix was produced starting from the ion level identifications with directLFQ (*67*) and LFQ intensities were averaged among the three technical replicates, $log_2$ transformed and filtered, retaining only hits identified in minimum 70 % of the samples in at least one condition for subsequent statistical analyses (n = 2307). To account for the large number of Missing Not At Random values in the differential expression analysis, the dataset was imputed with the MinProb function from the MSnbase R package (*68*), using the default *q* value of 0.01. Differential expression analysis was performed with the DEqMS package in R (*53*, *69*). Proteins were considered significantly differentially expressed if they displayed a $log_2$ fold change higher than 1.5, with an adjusted p-value < 0.05. Globally, 606 significant differentially expressed proteins (26 %) were identified across all conditions.

#### Supplementary Text

#### Trace metals depletion effect on *E. coli* aerobic growth

Similarly to anaerobic growth, fully aerobic cultures grown in M9 TM-rich and TM-depleted media exhibited a 35 % difference in growth yield between the two conditions (Fig. S5). The addition of copper and iron at 12 hours in the TM-depleted aerobic condition restored biomass yields to TM-rich levels (Fig. S5). While molybdenum and iron are critical for anaerobic growth (see main text), copper and iron play a fundamental role in aerobic metabolism.

Aerobic respiration depends on three terminal oxidases: quinol oxidases bo3, bd-I, and bd-II (see Table S3). The quinol oxidase bo3 is a heme-copper oxidase requiring both iron and copper. It is encoded by the cyoABCDE operon and expressed under high oxygen conditions (*70*). In contrast, quinol oxidases bd-I and bd-II, encoded by the cydAB and appBC operons, are heme oxidases that rely only on iron and operate under oxygen-limiting conditions. In fully aerobic growth, quinol oxidase bo3 is therefore expected to play a dominant role, highlighting the importance of iron and copper in supporting maximal aerobic respiration and growth.

##### Protein interaction network analysis of differentially expressed proteins

To highlight the physiological differences between cells grown anaerobically in M9 TM-rich and TM-depleted media, differentially expressed proteins were analyzed using STRING (*71*). The protein-protein interaction network was clustered into subnetworks of closely interacting proteins and each cluster was subjected to enrichment analysis using as a background the whole proteome. At each time point, distinct protein clusters were identified in the two conditions. By 6 h, the primary upregulated protein network in the TM-depleted condition consisted of proteins involved in iron uptake (Ton, Fhu, Fep) and in the biosynthesis of the siderophore enterobactin (Ent) (Fig. S9A). Additionally, subunits of the Suf system of the Fe-S cluster assembly were overexpressed. These proteins are involved in the Fe-S cluster biogenesis under iron limitation or oxidative stress, and may facilitate iron uptake from extracellular chelators. A group of ABC transporters involved in divalent cation import (Mnt, Zup, Zin, Znu) was also upregulated (Fig. S9A). Collectively, the upregulation of these protein networks highlights the metabolic effort for the acquisition of metals and describes a general cellular stress condition.

In contrast, during early growth in the TM-rich condition, the sole presence of TMs is associated with a clear activation of the anaerobic metabolic pathways: the two main upregulated protein clusters are related to the TCA cycle and oxidative phosphorylation (Fig. S9B). A third large cluster contains proteins involved in the assembly of metal cofactors (Moa for molybdopterin and Isc for Fe-S cluster). Other clusters upregulated in this condition included proteins involved in cell wall biosynthesis, nucleotide and amino acid metabolisms, and monosaccharide and oligopeptide transport. The overexpression of these networks indicate a physiological active replication phase in TM-rich condition, which appears instead impaired in the TM-depleted condition (Fig. S9B).

Similar patterns are observed after 12 (Fig. S10) and 24 h (Fig. S11). In the TM-depleted conditions, multiple interacting protein networks related to iron and transition metal acquisition and transport remain upregulated, indicating persistent TMs starvation (Fig. S10A and Fig. S11A). Interestingly, proteins involved in the biosynthesis of aromatic amino acids (His, Trp, Phe, Tyr) become increasingly overrepresented. A similar response has been observed in plants, where these amino acids contribute to protein synthesis and stress-related metabolites production, possibly highlighting a role of aromatic amino acids in stress adaptation (*74*). In contrast, in the TM-rich condition the bacterium continues to express proteins linked to active anaerobic metabolism and nitrate respiration (Nar, Frd, Nuo and Fdn) (Fig. S10B and Fig. S11B). The appearance of hydrogenases (Hyc, Hyf) and associated proteins, along with enzymes involved in oxygen consumption and the transition to fermentation, such as the GlpABC complex, is also more efficient compared to the TM-depleted cultures. The upregulation of the amino acids biosynthetic pathways suggests a physiological response to amino acids starvation in M9 minimal medium, while the upregulation of the protein networks related to cell replication indicates ongoing cellular growth and division in the TM-rich condition. This contrasts with the TM-depleted condition, where growth and replication are inhibited, in line with the experimental observations.

### Supplementary Figures

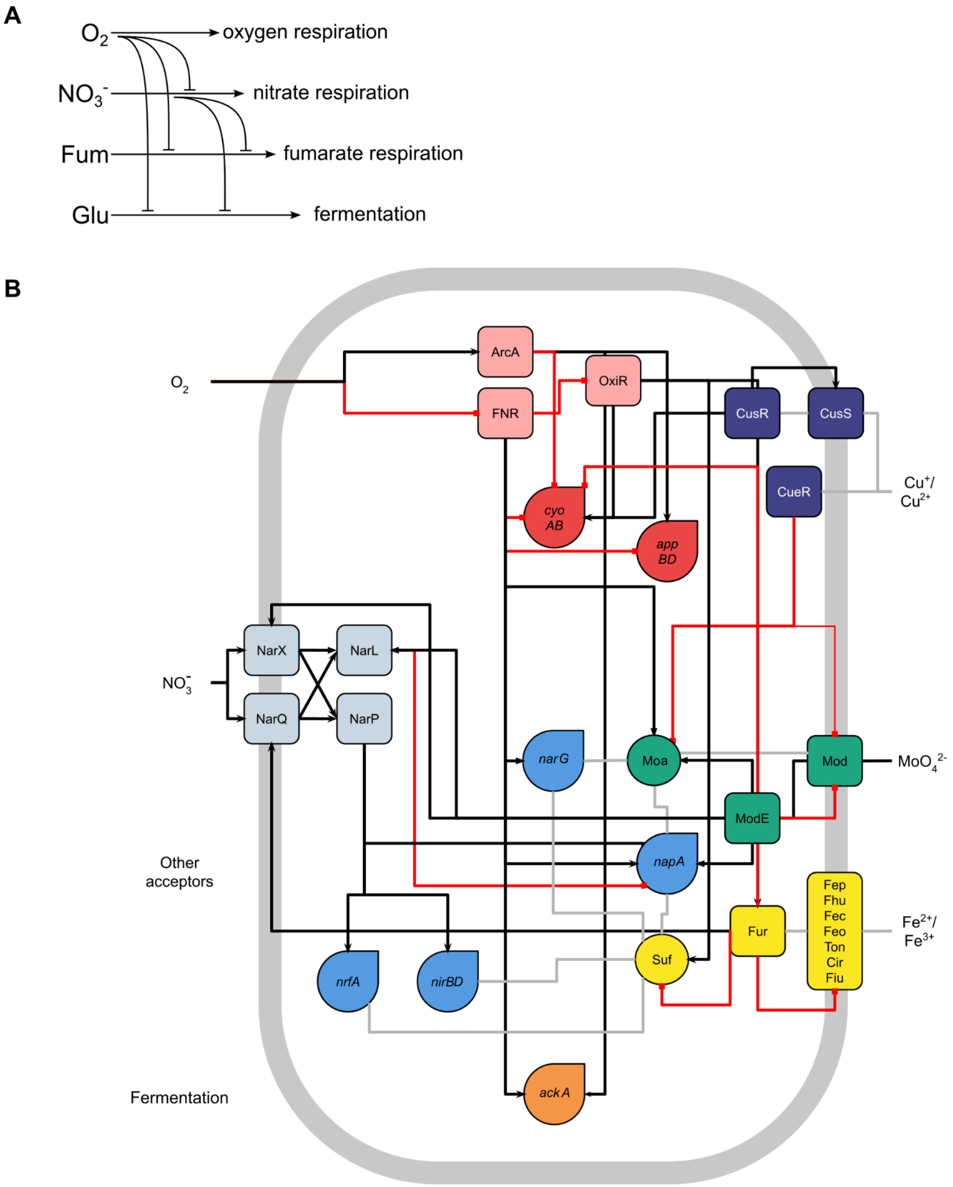


**Fig. S1.** Simplified summary of *E. coli* regulatory network for respiration and fermentation. (A) Regulation cascade of alternative respiratory pathways and fermentation. (B) Simplified schematics of the regulatory network controlling the switch from aerobic respiration, to nitrate respiration to fermentation and the associated metal cofactor regulators. Black arrows show positive regulations while red links with a square show negative regulation. Adapted from (*23*) and from RegulonDB (*72*).

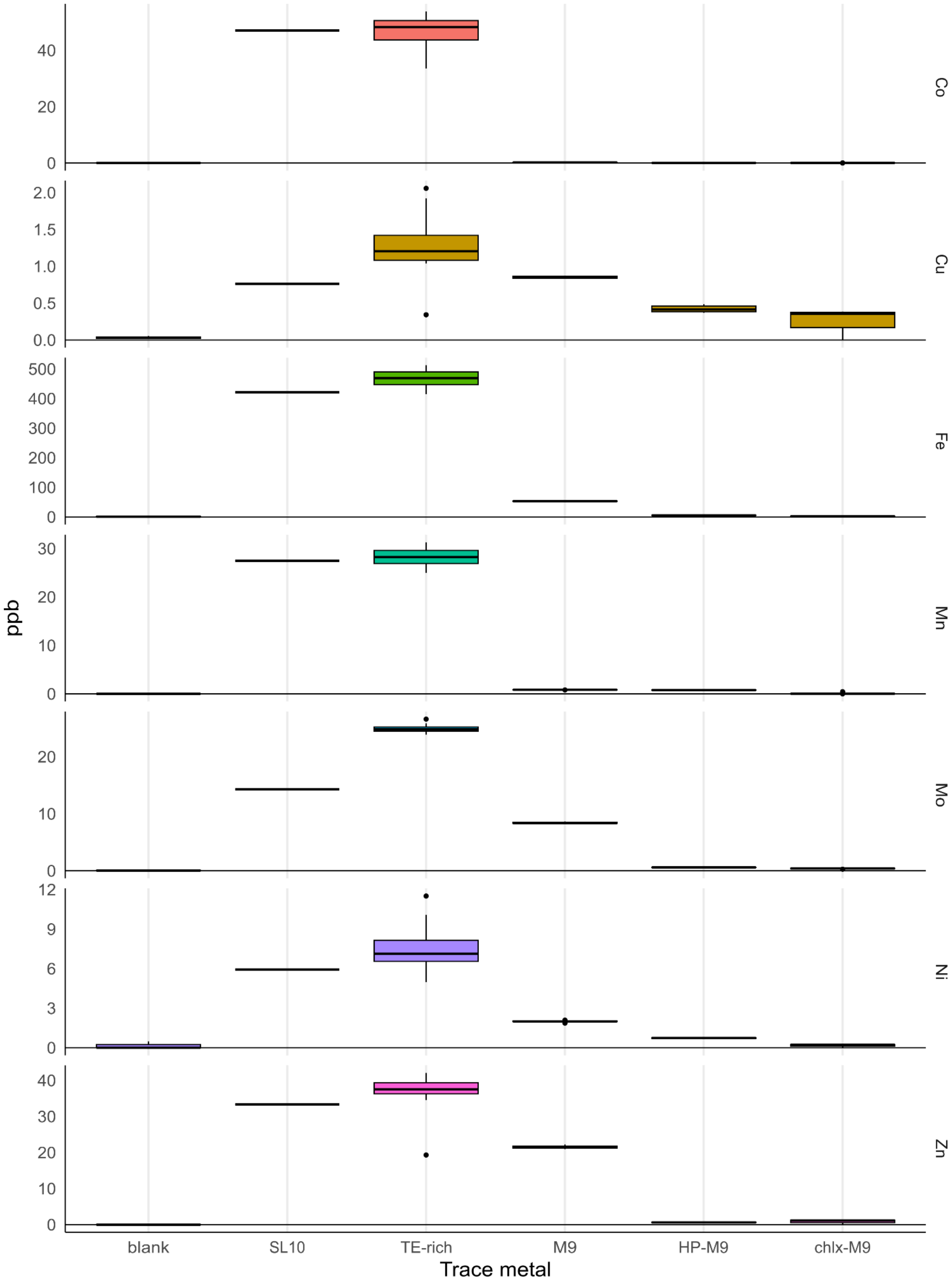


**Fig. S2.** Trace metal concentrations in the *E. coli* test media. Values (in ppb) for the laboratory blanks (blank) and the theoretical concentrations in pure M9 added with SL10 trace metal solution (SL10) are reported along with the values obtained for the TM-rich, standard M9, HP-M9 and chlx-M9 media.

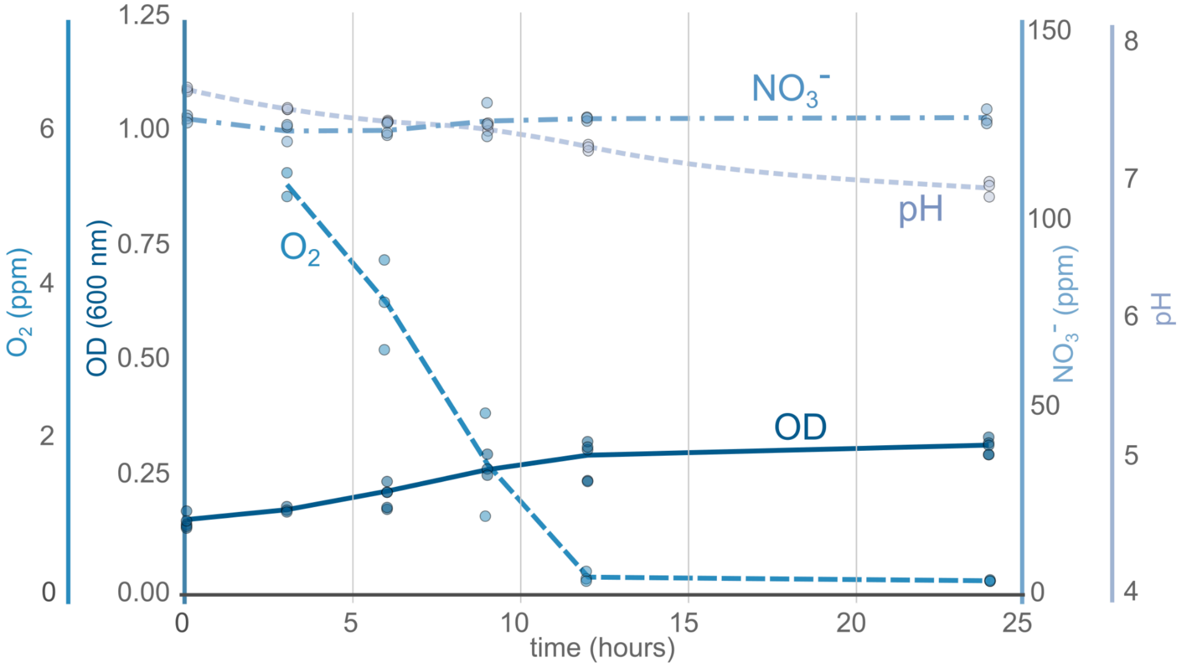


**Fig. S3.** *E. coli* anaerobic growth in chlx-M9 TM-depleted conditions, oxygen and nitrate concentrations and pH levels.

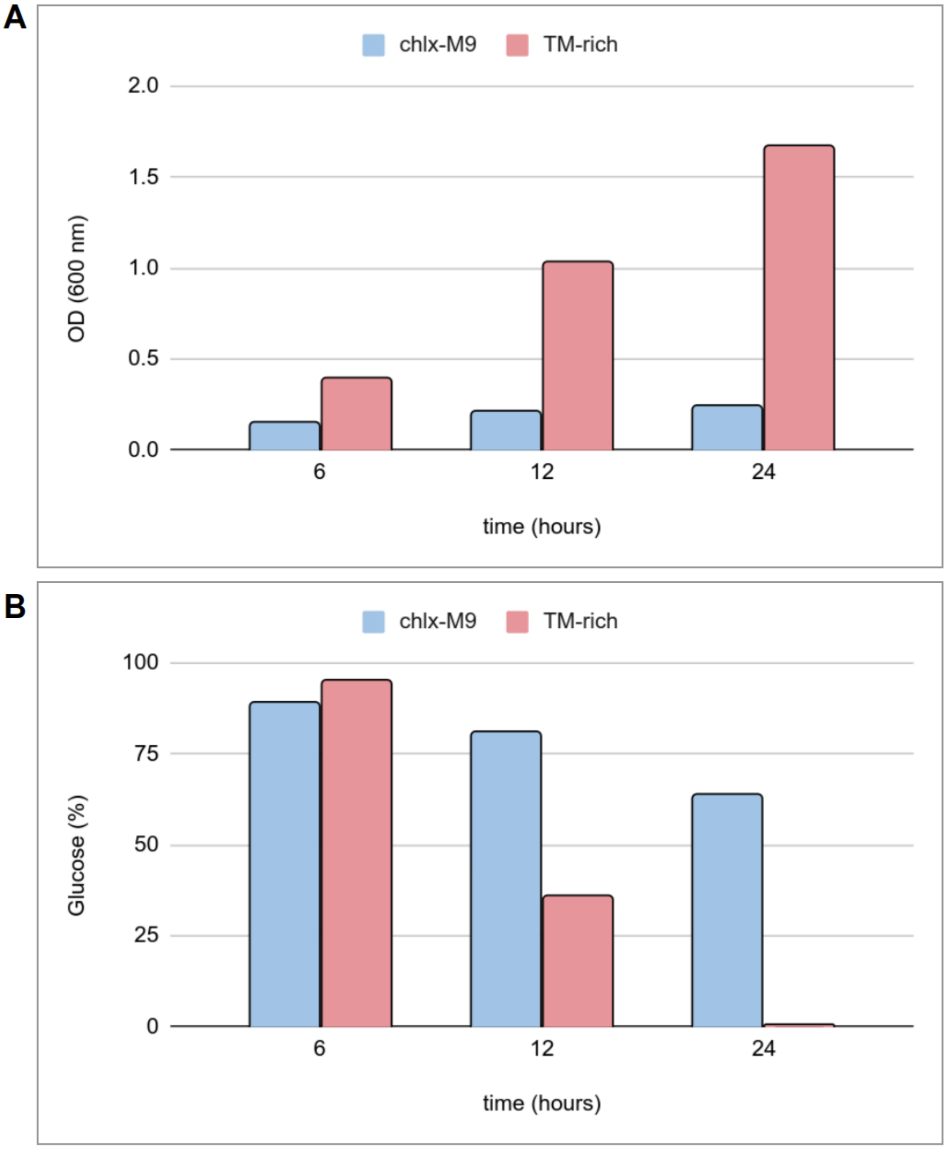


**Fig. S4.** *E. coli* aerobic growth and in TM-rich (red) and chlx-M9 TM-depleted (blue) conditions. Average values of optical densities (A) and residual glucose percentages (B) at selected time points (6, 12 and 24 h).

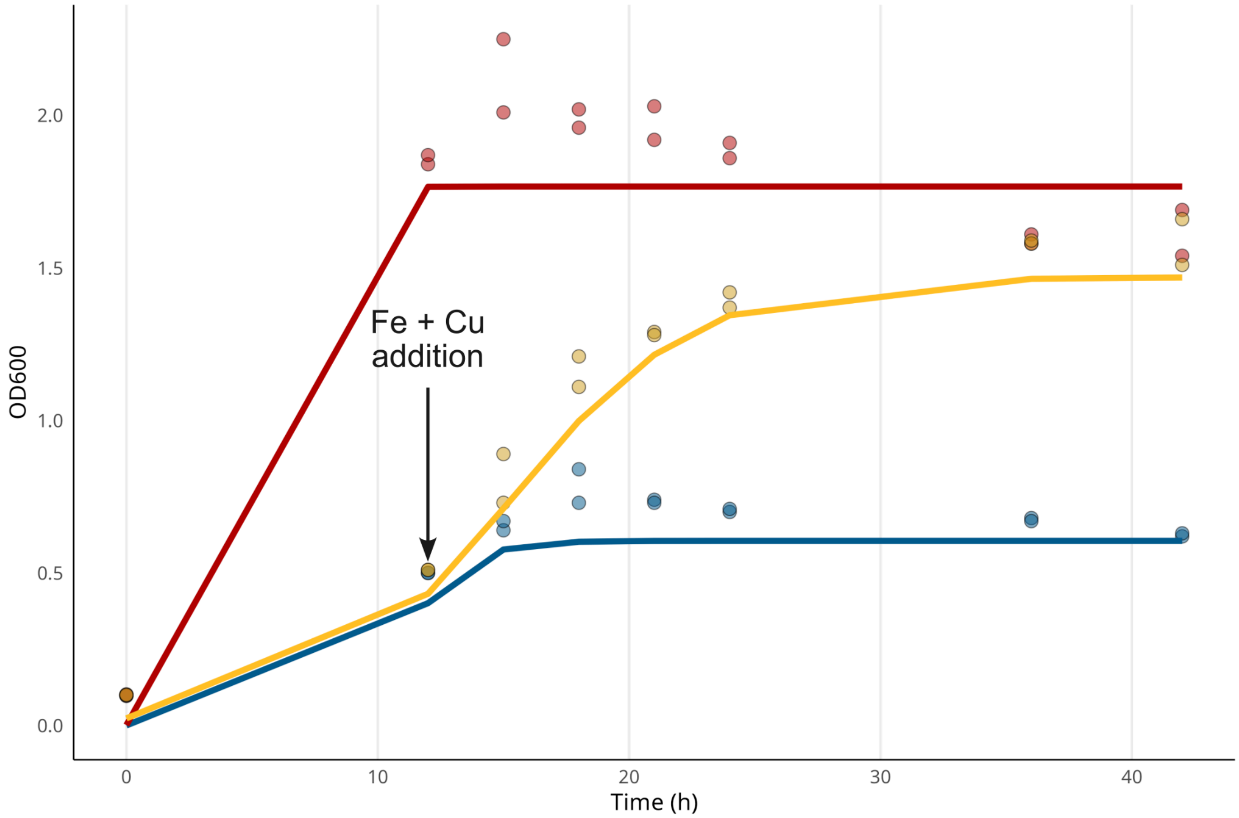


**Fig. S5.** The addition of copper and iron at 12 hours during aerobic growth restores growth to TM-rich levels. TM-rich growth (in red), TM-depleted HP-M9 with copper and iron supplemented at 12 hours (yellow) and TM-depleted HP-M9 growth (blue).

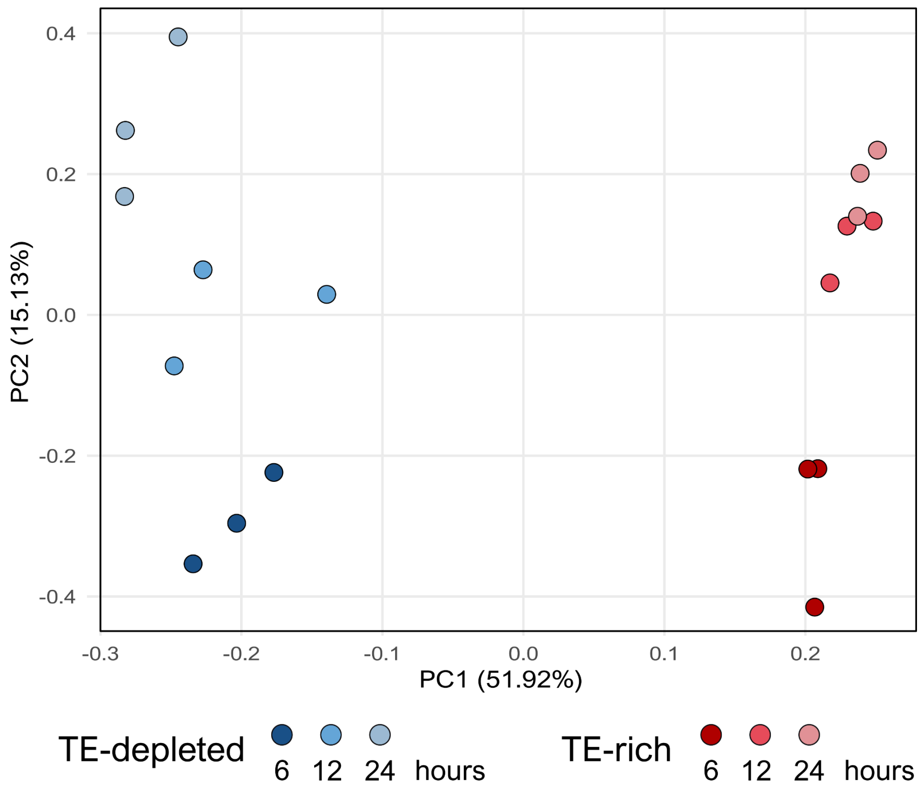


**Fig. S6.** Principal component analysis of the unimputed dataset obtained from the label free quantification proteomics experiments. Three biological replicates were analyzed for each condition, each resulting from the averaging of three technical replicates.

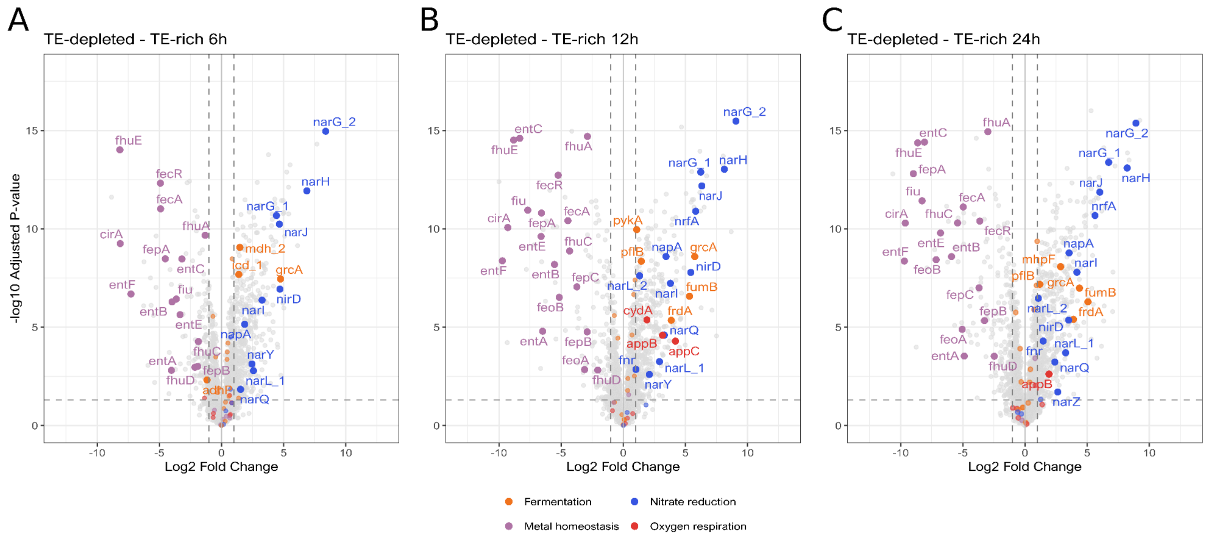


**Fig. S7.** Volcano plots highlighting differentially expressed proteins between the M9 TM-depleted (left) and TM-rich (right) cultures at 6 (A), 12 (B) and 24 h (C).

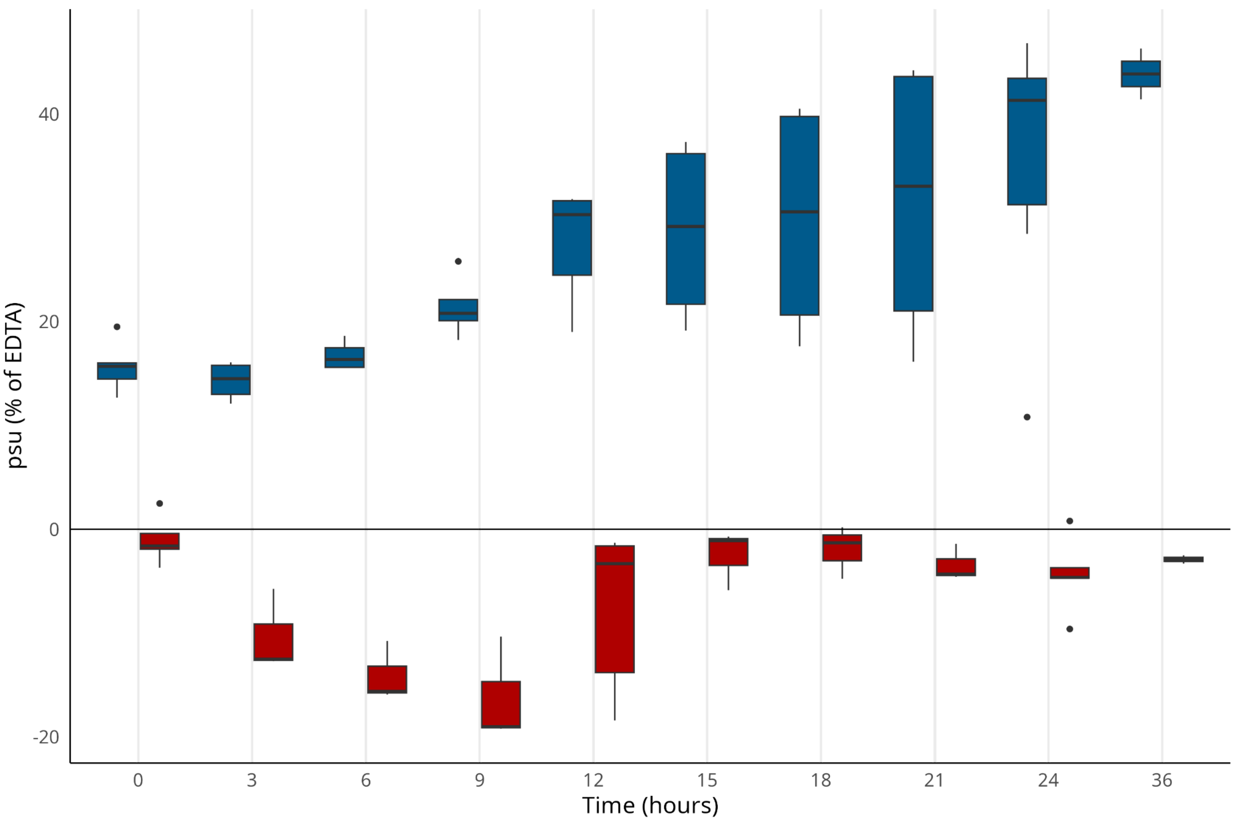


**Fig. S8.** Siderophores production during anaerobic growth of *E. coli* in M9 TM-depleted (blue) and TM-rich (red) conditions. Production was measured at selected time points using CAS assay and expressed as a siderophore production unit (psu) normalized to the positive control (EDTA).

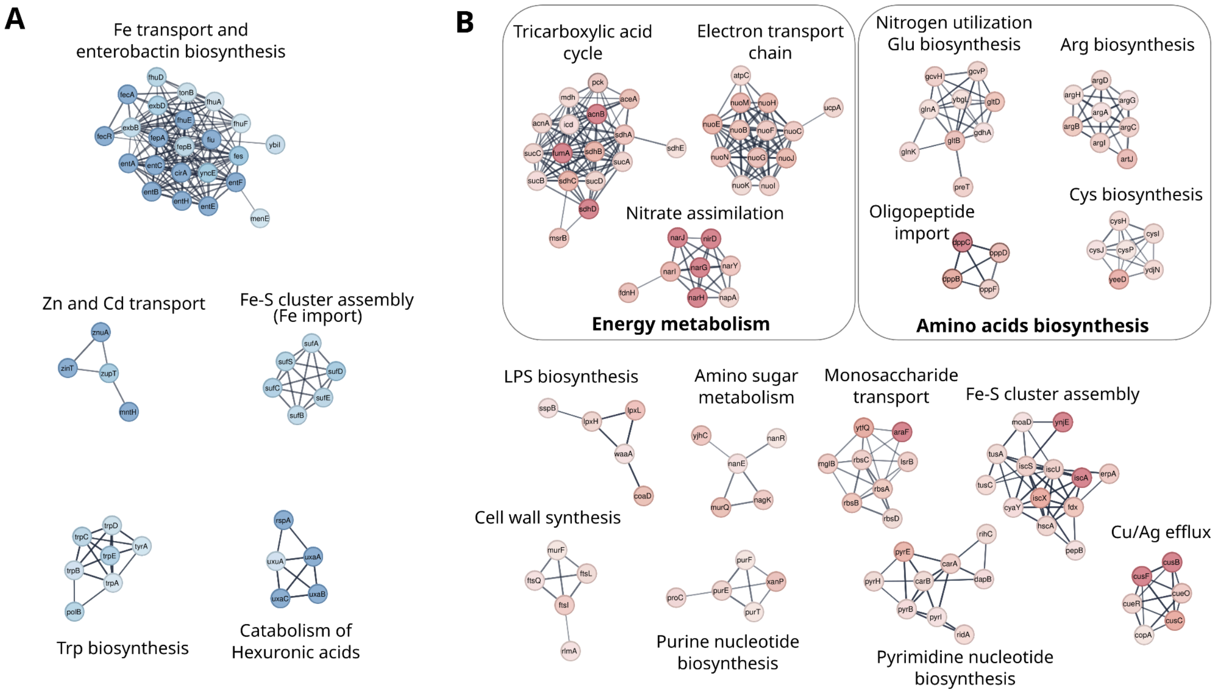


ref

**Fig. S9.** STRING analysis of differentially expressed proteins after 6 h of anaerobic growth of *E. coli* in M9 TM-depleted (A) and TM-rich (B) conditions. Clusters of proteins involved in the same cellular functions are represented in subgroups.

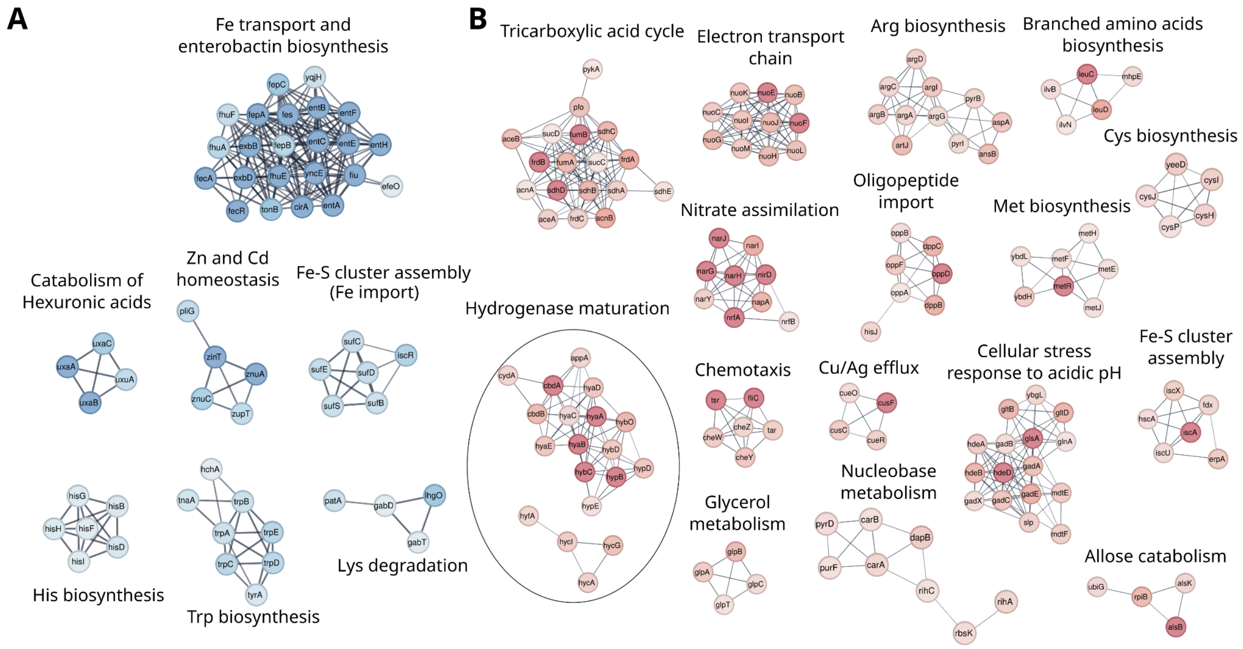


**Fig. S10.** STRING analysis of differentially expressed proteins after 12 h of anaerobic growth of *E. coli* in M9 TM-depleted (A) and TM-rich (B) conditions. Clusters of proteins involved in the same cellular functions are represented in subgroups.

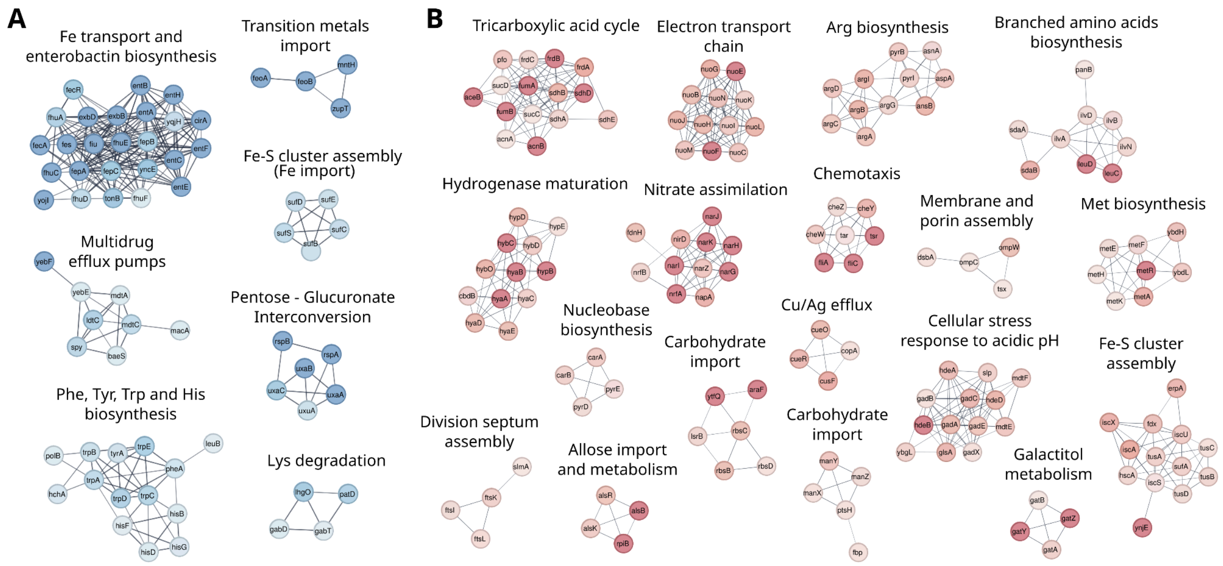


**Fig. S11.** STRING analysis of differentially expressed proteins after 24 h of anaerobic growth of *E. coli* in M9 TM-depleted (A) and TM-rich (B) conditions.

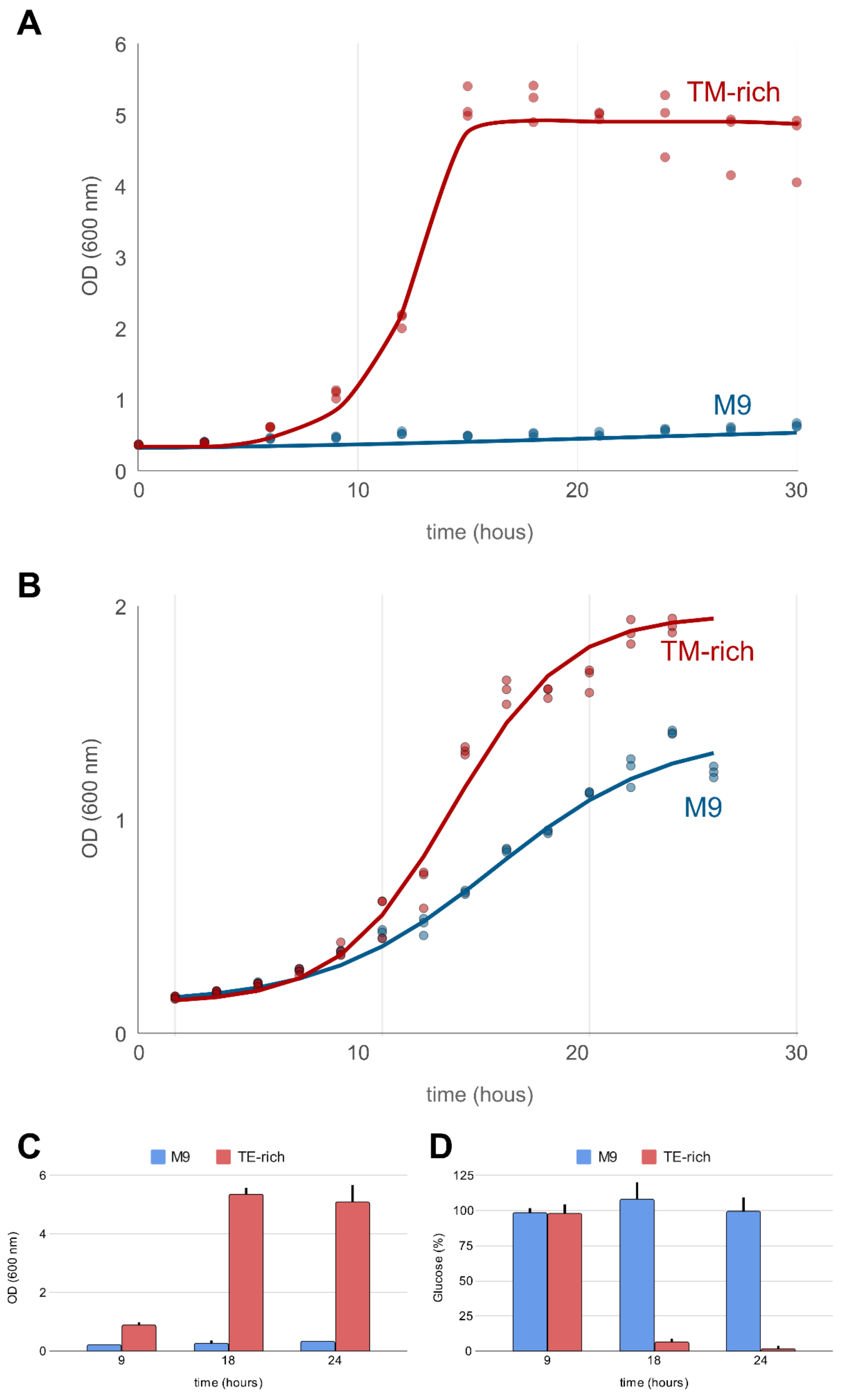


**Fig. S12.** *Bacillus subtilis* aerobic (A) and anaerobic (B) growth in M9 TM-depleted (blue) and TM-rich (red) conditions. Average values of optical densities (C) and residual glucose percentages (D) at selected time points (6, 12 and 24 h).

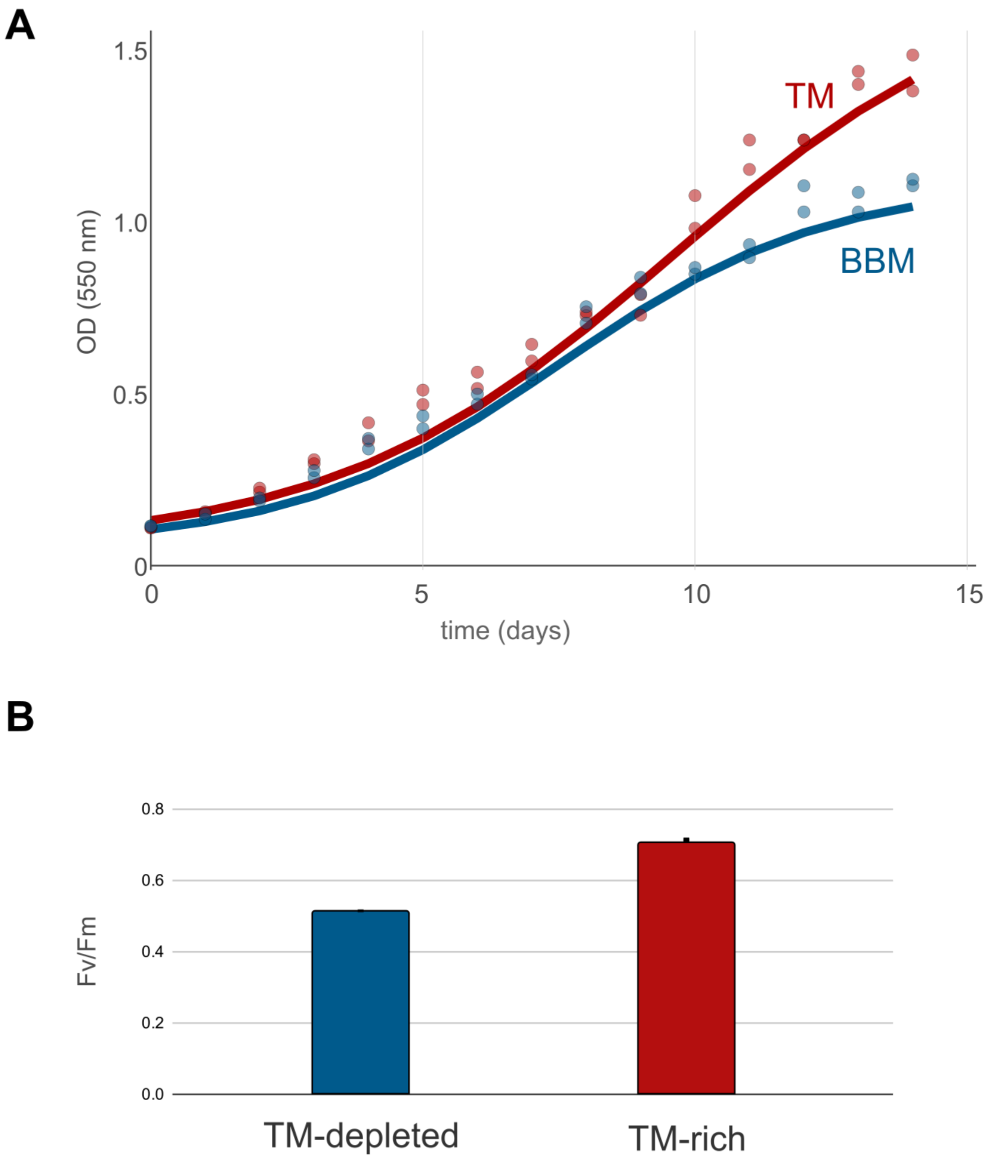


**Fig. S13.** *Stichococcus bacillaris* Nägeli growth in M9 TM-depleted (blue) and TM-rich (red) conditions (A). Maximum quantum efficiency of PSII photochemistry has been measured in both conditions (B).

#### Supplementary Tables

**Table S1.** Diversity of electron acceptors and metals cofactor used by terminal reductases in MAGs from different environments. MAGs downloaded from the JGI GEMs Genomic catalogue of Earth's microbiomes (*12*).

| Environment | n. MAGs | MAGs with AEA[a] | MAGs TR[b] metal cofactor |
|---|---|---|---|
| Aquatic | 19,300 | 85.4 % | 74.0 % |
| Human | 16,441 | 62.3 % | 23.8 % |
| Terrestrial | 3,411 | 89.6 % | 80.6 % |
| Built environment | 2,640 | 87.7 % | 84.7 % |
| Other environments | 8,096 | 80.1 % | 48.9 % |
| **Total GEM collection** | **52,515** | **78.1 %** | **70.6 %** |

a. Alternative electron acceptors.
b. Terminal reductases.

**Table S2**. Terminal reductases and HMM models used for the annotation of the MAGs of the GEM collection.

| Enzyme full name | Enzyme short name | Metal cofactors | Pathway | HMM acc. number |
|---|---|---|---|---|
| Thiosulfate reductase | PhsA | Mo, Fe | Thiosulfate reduction | (*73*) |
| Dissimilatory sulfite reductase | DsrA | Fe | Dissimilatory sulfate reduction | TIGR02064 |
| Periplasmic nitrate reductase | NapA | Mo, Fe | Nitrate reduction | TIGR01706 |
| Membrane bound nitrate reductase | NarG | Mo, Fe | Nitrate reduction | PF02665 |
| Cytochrome c oxidase | CyoB | Cu, Fe | Oxygen respiration, aerobic | PF00115 |
| Cytochrome d ubiquinol oxidase | CydA | Fe | Oxygen respiration, microaerophilic | PF01654 |

**Table S3.** Metabolic diversity of *E. coli* and metal containing oxidoreductases and key metal containing enzymes for each pathway

| Conditions | Metabolis | Pathway | Redox couple | Enzyme name | Gene names | EC number | PDB accession | Metals | Metal cofactors |
|---|---|---|---|---|---|---|---|---|---|
| Aerobic | Aerobic respiration | Oxygen reduction | $O_2/H_2O$ | Quinol oxidase bo3 | *cyoABCDE* | 7.1.1.3 | 8GO3 | Cu, Fe | Fe (heme O), Fe (protoporphyrin IX), $Cu^{2+}$ (ion) |
| Microaerophilic | | | | Quinol oxidase bd-I | *cydAB* | 7.1.1.7 | 6RKO | Fe | Fe (heme b/c), Fe (cis-heme D) |
| | | | | Quinol oxidase bd-II | *appBC (=cyxAB)* | 7.1.1.7 | 7OSE | Fe | Fe (heme b/c), Fe (cis-heme D) |
| Anaerobic | Denitrification | Nitrate reduction | $NO_3^-/NO_2^-$ | Membrane bound nitrate reductase A | *narGHJI* | 1.7.5.1 | 1Q16 | Mo, Fe | $Mo^{6+}$ (ion within molybdopterin dinucleotide MD1), Fe (protoporphyrin IX), Fe (4Fe-4S), Fe (3Fe-4S) |
| | | | | Nitrate reductase Z | *narZYWV* | 1.7.5.1 | -[a] | Mo, Fe | $Mo^{6+}$ (ion within molybdopterin dinucleotide MD1), Fe (protoporphyrin IX), Fe (4Fe-4S), Fe (3Fe-4S) |
| | | | | Nitrate reductase, periplasmic | *napFDAGHBC* | 1.9.6.1 | 3ML1[b] | Mo, Fe | $Mo^{6+}$ (dioxothiomolybdenum ion within bis-molybdopterin dinucleotide MGD), Fe (heme c), Fe (4Fe-4S) |
| | | Nitrite reduction | $NO_2^-/NH_4^+$ | Nitrite reductase | *nrfABCDEFG* | 1.7.2.2 | 2J7A[c] | Fe | Fe (heme c) |
| | | | | Nitrite reductase (NADH) | *nirBD* | 1.7.1.15 | AF-P08201-F1, 2JO6 | Fe | Fe (2Fe-2S rieske), Fe (4Fe-4S), Fe (siroheme) |
| | Other anaerobic respiratory pathways | DMSO reduction | DMSO/DMS | DMSO reductase | *dmsABC* | 1.8.5.3 | 1DMS | Mo, Fe | $Mo^{6+}$ (ion with bis-molybdopterin dinucleotide MGD), Fe (4Fe-4S), Fe (3Fe-4S) |
| | | TMAO reduction | TMAO/TMA | TMAO reductase | *torCAD* | 1.7.2.3 | 1TMO[d] | Mo, Fe | $Mo^{6+}$ (ion with bis-molybdopterin dinucleotide MGD) |
| | | Fumarate reduction | fumarate/ succinate | Fumarate reductase | *frdABCD* | 1.3.1.6 | 1KF6 | Fe | Fe (4Fe-4S), Fe (3Fe-4S), Fe (2Fe-2S) |
| | Fermentation | Fermentation | acetaldehyde/ ethanol | alcohol dehydrogenase | *adh* | 1.1.1.1 | 6TQM | Fe | $Fe^{3+}$ (ion) |
| | | | fumarate/ succinate | succinate dehydrogenase | *sdh* | 1.3.5.1 | 5XMJ[e] | Fe | Fe (protoporphyrin IX), Fe (4Fe-4S), Fe (3Fe-4S), Fe (2Fe-2S) |
| | | | $formate/CO_2$ | formate hydrogenlyase | *hyc* | 1.12.7.2 | 7Z0S | Ni, Fe | Ni (NiFe), Fe (4Fe-4S), Fe (carbonmonoxide-dicyano iron) |
| | | | | | *fdhF* | 1.17.99.7 | 1FDO | Mo, Fe | $Mo^{6+}$ (ion with bis-molybdopterin dinucleotide MGD), Fe (4Fe-4S) |
| | | | $formate/CO_2$ | formate dehydrogenase | *fdnGHI* | 1.17.5.3 | 1KQF | Mo, Fe | $Mo^{6+}$ (ion with bis-molybdopterin dinucleotide MGD), Fe (4Fe-4S), Fe (protoporphyrin IX) |
| | | | pyruvate/ acetate | Pyruvate dehydrogenase | *poxB* | 1.2.5.1 | 3EYA | Mg | $Mg^{2+}$ (ion) |

a. The PDB structure for *E. coli* is missing. Based on sequence similarity it is expected to be similar to NarG.
b. The PDB structure for *E. coli* is missing. The structure for *Cupriavidus necator* is reported.
c. The complete PDB structure for *Desulfovibrio vulgaris* is reported.
d. The PDB structure for *E. coli* is missing. The structure for *Shewanella massilia* is reported.
e. The PDB structure for *E. coli* is missing. The structure for *Desulfovibrio gigas* is reported.

**Table S4**. Concentration of the trace metals of biological interest in the *E. coli* growth media.

| condition | media | trace | treatment | Mn (ppb) | Fe (ppb) | Co (ppb) | Ni (ppb) | Cu (ppb) | Zn (ppb) | Mo (ppb) |
|---|---|---|---|---|---|---|---|---|---|---|
| theoretical | SL10 | SL10 | theoretical | 27.47 | 421.07 | 47.06 | 5.93 | 0.76 | 33.34 | 14.30 |
| rich | M9 | SL10 | standard salts | 0.84 ± 0.02 | 53.14 ± 1.27 | 0.22 ± 0.00 | 1.99 ± 0.07 | 0.85 ± 0.02 | 21.53 ± 0.46 | 8.42 ± 0.15 |
| depleted | HP-M9 | none | high-purity salts | 0.75 ± 0.03 | 5.10 ± 0.14 | 0.02 ± 0.00 | 0.74 ± 0.04 | 0.42 ± 0.05 | 0.62 ± 0.07 | 0.6 ± 0.02 |
| | chlx-M9 | | high-purity salts, chelex treated | 0.04 ± 0.01 | 2.2 ± 0.04 | 0.06 ± 0.00 | 0.23 ± 0.01 | 0.36 ± 0.02 | 1.28 ± 0.16 | 0.41 ± 0.01 |

**Table S5.** Unique proteins associated with the main metabolic processes under investigation identified at the three time points in only one culture condition (TM-depleted or TM-rich).

| Gene | Protein | Only found in | Time | Process |
|---|---|---|---|---|
| entD | 242384E_coliDH5_03564 | TM-depleted | 6 h | Metal Homeostasis |
| fecA | 242384E_coliDH5_02531 | TM-depleted | 6 h | Metal Homeostasis |
| fecR | 242384E_coliDH5_02530 | TM-depleted | 6 h | Metal Homeostasis |
| fhuD | 242384E_coliDH5_01275 | TM-depleted | 6 h | Metal Homeostasis |
| fhuE | 242384E_coliDH5_03238 | TM-depleted | 6 h | Metal Homeostasis |
| fhuF | 242384E_coliDH5_02448 | TM-depleted | 6 h | Metal Homeostasis |
| fumB | 242384E_coliDH5_00834 | TM-rich | 6 h | Fermentation |
| tdcE | 242384E_coliDH5_01953 | TM-rich | 6 h | Fermentation |
| napA | 242384E_coliDH5_00952 | TM-rich | 6 h | Nitrate reduction |
| narG_1 | 242384E_coliDH5_02223 | TM-rich | 6 h | Nitrate reduction |
| narG_2 | 242384E_coliDH5_02224 | TM-rich | 6 h | Nitrate reduction |
| narH | 242384E_coliDH5_02222 | TM-rich | 6 h | Nitrate reduction |
| narI | 242384E_coliDH5_02220 | TM-rich | 6 h | Nitrate reduction |
| narJ | 242384E_coliDH5_02221 | TM-rich | 6 h | Nitrate reduction |
| narU | 242384E_coliDH5_00381 | TM-rich | 6 h | Nitrate reduction |
| narY | 242384E_coliDH5_00379 | TM-rich | 6 h | Nitrate reduction |
| nirC | 242384E_coliDH5_01602 | TM-rich | 6 h | Nitrate reduction |
| nirD | 242384E_coliDH5_01601 | TM-rich | 6 h | Nitrate reduction |
| nrfA | 242384E_coliDH5_00888 | TM-rich | 6 h | Nitrate reduction |
| cyoD | 242384E_coliDH5_02710 | TM-rich | 6 h | Oxygen respiration |
| entD | 242384E_coliDH5_03564 | TM-depleted | 12 h | Metal Homeostasis |
| entF | 242384E_coliDH5_03560 | TM-depleted | 12 h | Metal Homeostasis |
| fecA | 242384E_coliDH5_02531 | TM-depleted | 12 h | Metal Homeostasis |
| fecR | 242384E_coliDH5_02530 | TM-depleted | 12 h | Metal Homeostasis |
| feoA | 242384E_coliDH5_01642 | TM-depleted | 12 h | Metal Homeostasis |
| fepD | 242384E_coliDH5_03556 | TM-depleted | 12 h | Metal Homeostasis |
| fhuB | 242384E_coliDH5_01276 | TM-depleted | 12 h | Metal Homeostasis |
| fhuD | 242384E_coliDH5_01275 | TM-depleted | 12 h | Metal Homeostasis |
| fhuE | 242384E_coliDH5_03238 | TM-depleted | 12 h | Metal Homeostasis |
| fhuF | 242384E_coliDH5_02448 | TM-depleted | 12 h | Metal Homeostasis |
| fieF | 242384E_coliDH5_02839 | TM-depleted | 12 h | Metal Homeostasis |
| fsr | 242384E_coliDH5_02759 | TM-depleted | 12 h | Metal Homeostasis |
| adhP | 242384E_coliDH5_00391 | TM-rich | 12 h | Fermentation |
| fumB | 242384E_coliDH5_00834 | TM-rich | 12 h | Fermentation |
| tdcE | 242384E_coliDH5_01953 | TM-rich | 12 h | Fermentation |
| napA | 242384E_coliDH5_00952 | TM-rich | 12 h | Nitrate reduction |
| narG_1 | 242384E_coliDH5_02223 | TM-rich | 12 h | Nitrate reduction |
| narG_2 | 242384E_coliDH5_02224 | TM-rich | 12 h | Nitrate reduction |
| narH | 242384E_coliDH5_02222 | TM-rich | 12 h | Nitrate reduction |
| narI | 242384E_coliDH5_02220 | TM-rich | 12 h | Nitrate reduction |
| narJ | 242384E_coliDH5_02221 | TM-rich | 12 h | Nitrate reduction |
| narU | 242384E_coliDH5_00381 | TM-rich | 12 h | Nitrate reduction |
| nirC | 242384E_coliDH5_01602 | TM-rich | 12 h | Nitrate reduction |

| | | | | |
|---|---|---|---|---|
| nrfA | 242384E_coliDH5_00888 | TM-rich | 12 h | Nitrate reduction |
| nrfB | 242384E_coliDH5_00887 | TM-rich | 12 h | Nitrate reduction |
| appB | 242384E_coliDH5_00321 | TM-rich | 12 h | Oxygen respiration |
| cirA | 242384E_coliDH5_03045 | TM-depleted | 24 h | Metal Homeostasis |
| entC | 242384E_coliDH5_03553 | TM-depleted | 24 h | Metal Homeostasis |
| entD | 242384E_coliDH5_03564 | TM-depleted | 24 h | Metal Homeostasis |
| entF | 242384E_coliDH5_03560 | TM-depleted | 24 h | Metal Homeostasis |
| fecA | 242384E_coliDH5_02531 | TM-depleted | 24 h | Metal Homeostasis |
| fecR | 242384E_coliDH5_02530 | TM-depleted | 24 h | Metal Homeostasis |
| feoA | 242384E_coliDH5_01642 | TM-depleted | 24 h | Metal Homeostasis |
| feoB | 242384E_coliDH5_01643 | TM-depleted | 24 h | Metal Homeostasis |
| fepC | 242384E_coliDH5_03558 | TM-depleted | 24 h | Metal Homeostasis |
| fepD | 242384E_coliDH5_03556 | TM-depleted | 24 h | Metal Homeostasis |
| fes | 242384E_coliDH5_03562 | TM-depleted | 24 h | Metal Homeostasis |
| fhuB | 242384E_coliDH5_01276 | TM-depleted | 24 h | Metal Homeostasis |
| fhuC | 242384E_coliDH5_01274 | TM-depleted | 24 h | Metal Homeostasis |
| fhuD | 242384E_coliDH5_01275 | TM-depleted | 24 h | Metal Homeostasis |
| fhuE | 242384E_coliDH5_03238 | TM-depleted | 24 h | Metal Homeostasis |
| fhuF | 242384E_coliDH5_02448 | TM-depleted | 24 h | Metal Homeostasis |
| fieF | 242384E_coliDH5_02839 | TM-depleted | 24 h | Metal Homeostasis |
| fiu | 242384E_coliDH5_00147 | TM-depleted | 24 h | Metal Homeostasis |
| fumB | 242384E_coliDH5_00834 | TM-rich | 24 h | Fermentation |
| tdcE | 242384E_coliDH5_01953 | TM-rich | 24 h | Fermentation |
| napA | 242384E_coliDH5_00952 | TM-rich | 24 h | Nitrate reduction |
| narG_1 | 242384E_coliDH5_02223 | TM-rich | 24 h | Nitrate reduction |
| narG_2 | 242384E_coliDH5_02224 | TM-rich | 24 h | Nitrate reduction |
| narH | 242384E_coliDH5_02222 | TM-rich | 24 h | Nitrate reduction |
| narI | 242384E_coliDH5_02220 | TM-rich | 24 h | Nitrate reduction |
| narJ | 242384E_coliDH5_02221 | TM-rich | 24 h | Nitrate reduction |
| narU | 242384E_coliDH5_00381 | TM-rich | 24 h | Nitrate reduction |
| nrfA | 242384E_coliDH5_00888 | TM-rich | 24 h | Nitrate reduction |
| nrfB | 242384E_coliDH5_00887 | TM-rich | 24 h | Nitrate reduction |

**Table S6.** TY medium composition.

| Reagent | Concentration [g/L] |
|---|---|
| Tryptone | 10 |
| Yeast extract | 5 |
| NaCl | 10 |

**Table S7.** M9 medium composition.

| Reagent | Concentration [g/L] | Molarity [mM] |
|---|---|---|
| $Na_2HPO_4$ | 6 | 42.26 |
| $KH_2PO_4$ | 3 | 22.05 |
| $NH_4Cl$ | 1 | 18.69 |
| NaCl | 0.5 | 8.55 |
| $MgSO_4$ | 0.24 | 2 |
| $CaCl_2$ | 0.01 | 0.1 |
| $KNO_3$ | 0.20 | 2 |
| Thiamine-HCl ● $2H_2O$ | 0.001 | 0.003 |
| Glucose | 2 | 11.10 |

**Table S8.** SL10 trace element composition.

| Reagent | Concentration [µg/L] | Molarity [µM] |
|---|---|---|
| $FeCl_2 \bullet 4\ H_2O$ | 421.07 | 7.55 |
| $ZnCl_2$ | 33.34 | 0.51 |
| $MnCl_2 \bullet 4\ H_2O$ | 27.47 | 0.51 |
| $H_3BO_3$ | 1.027 | 0.10 |
| $CoCl_2 \bullet 6\ H_2O$ | 47.06 | 0.80 |
| $CuCl_2 \bullet 2\ H_2O$ | 0.76 | 0.01 |
| $NiCl_2 \bullet 6\ H_2O$ | 5.92 | 0.10 |
| $Na_2MoO_4 \bullet 2\ H_2O$ | 14.30 | 0.15 |

**Table S9**. Composition of the Bold's Basal Medium (BBM) used for *Stichococcus bacillaris* Nägeli growth experiments.

| Base reagent | Concentration [g/L] | Molarity [mM] |
|---|---|---|
| $NaNO_3$ | 0.250 | 2.940 |
| $CaCl_2 \bullet H_2O$ | 0.025 | 0.194 |
| $MgSO_4 \bullet 7\ H_2O$ | 0.075 | 0.305 |
| $K_2HPO_4$ | 0.075 | 0.431 |
| $KH_2PO_4$ | 0.175 | 1.286 |
| NaCl | 0.025 | 0.428 |
| EDTA | 0.050 | 0.154 |
| $H_3BO_3$ | 0.011 | 0.184 |
| $FeSO_4 \bullet 7\ H_2O$ | 0.008 | 0.027 |

**Table S10**. Composition of the Bold's Basal Medium (BBM) trace metal solution used for *Stichococcus bacillaris* Nägeli growth experiments.

| Trace metal solution | Concentration [g/L] | Molarity [mM] |
|---|---|---|
| $Na_2EDTA$ | 4.500 | 12.080 |
| $FeCl_3 \bullet 6\ H_2O$ | 0.582 | 2.150 |
| $MnCl_2 \bullet 4\ H_2O$ | 0.246 | 1.240 |
| $ZnCl_2$ | 0.030 | 0.123 |
| $CoCl_2$ | 0.012 | 0.050 |
| $Na_2MoO_4 \bullet 2\ H_2O$ | 0.024 | 0.099 |
